# METTL18-mediated histidine methylation on RPL3 modulates translation elongation for proteostasis maintenance

**DOI:** 10.1101/2021.07.29.454307

**Authors:** Eriko Matsuura-Suzuki, Tadahiro Shimazu, Mari Takahashi, Kaoru Kotoshiba, Takehiro Suzuki, Kazuhiro Kashiwagi, Yoshihiro Sohtome, Mai Akakabe, Mikiko Sodeoka, Naoshi Dohmae, Takuhiro Ito, Yoichi Shinkai, Shintaro Iwasaki

**Author notes:** These authors contributed equally (Eriko Matsuura-Suzuki, Tadahiro Shimazu). Corresponding authors Correspondence should be addressed to Tadahiro S., T.I., Yoichi S., and S.I., and (S.I.).

## Abstract

Protein methylation occurs predominantly on lysine and arginine residues, but histidine also serves as a substrate for the modification. However, a limited number of enzymes responsible for this modification have been reported. Moreover, the biological role of histidine methylation has remained poorly understood. Here, we report that human METTL18 is a histidine methyltransferase for the ribosomal protein RPL3 and that the modification specifically slows ribosome traverse on tyrosine codons, allowing the proper folding of synthesized proteins. By performing an *in vitro* methylation assay with a methyl donor analog and quantitative mass spectrometry, we found that His245 of RPL3 is methylated at the τ-*N* position by METTL18. Structural comparison of the modified and unmodified ribosomes showed stoichiometric modification and suggested a role in translation tuning. Indeed, genome-wide ribosome profiling revealed suppressed ribosomal translocation at tyrosine codons by RPL3 methylation. Because the slower elongation provides enough time for nascent protein folding, RPL3 methylation protects cells from the cellular aggregation of Tyr-rich proteins. Our results reveal histidine methylation as an example of a “ribosome code” that ensures proteome integrity in cells.

## Introduction

Contributing to the critical role of epigenetics, protein methylation is an integral posttranslational modification (PTM). This modification influences the function of proteins and provides a convertible platform for the modulation of cellular processes, including interactions with other molecules, protein structure, localization, and enzymatic activity (Biggar and Li, 2015; Clarke, 2018; Murn and Shi, 2017; Rodríguez-Paredes and Lyko, 2019). Whereas the majority of protein methylation has been observed on lysine and arginine amino acids (Biggar and Li, 2015; Clarke, 2018; Murn and Shi, 2017; Rodríguez-Paredes and Lyko, 2019), histidine residues also supply alternative target sites for methylation. Although this modification was previously thought to be restricted to limited proteins (Webb et al., 2010), a recent comprehensive survey suggested widespread occurrence in the proteome (Davydova et al., 2021; Ning et al., 2016; Wilkinson et al., 2019).

Two distinct nitrogen atoms in histidine could be methylated: the π*-N* position (π*-N-*methylhistidine or 1-methylhistidine) and the τ-*N* position (τ*-N-*methylhistidine or 3-methylhistidine) (Figure S1A). Regarding their responsible enzymes, methyltransferase-like (METTL) 9—a seven β-strand methyltransferase—and SET domain containing 3 (SETD3) are, as of yet, the only known mammalian methyltransferases for π*-N*- methylhistidine (Davydova et al., 2021) and τ*-N-*methylhistidine (Dai et al., 2019; Guo et al., 2019; Kwiatkowski et al., 2018; Wilkinson et al., 2019; Zheng et al., 2020), respectively. Nevertheless, the landscape of histidine methyltransferase-substrate pairs and, more importantly, the physiological functions of the modification have remained largely elusive.

Recent emerging studies have shown that the ribosome is a hotspot for PTM (Emmott et al., 2019; Simsek and Barna, 2017). This led to the concept of the “ribosome code”, analogous to the so-called “histone code”, in which PTMs on the macromolecule specify function and tune gene expression. Indeed, PTMs on ribosomal proteins may regulate protein synthesis in specific contexts, such as a subset of transcripts (Imami et al., 2018; Kapasi et al., 2007; Mazumder et al., 2003), the cell cycle (Imami et al., 2018), stress response (Higgins et al., 2015; Matsuki et al., 2020), and differentiation (Werner et al., 2015).

In this work, we studied the previously uncharacterized methyltransferase METTL18. Through a survey of the substrate, we found that this protein catalyzes τ*-N-* methylation on His245 of the ribosomal protein large subunit (RPL) 3 (uL3 as universal nomenclature). Cryo-electron microscopy (cryo-EM) suggested that τ*-N-*methylation interferes with the interaction of His245, which is located close to the peptidyl transferase center (PTC), with G1595 in the loop of helix 35 in 28S rRNA. Genome-wide ribosome profiling and quantitative proteome analysis revealed that the τ*-N*-methylhistidine on RPL3 retards translation elongation at Tyr codons, allows the proper folding of synthesized proteins, and thus ensures healthy proteostasis in cells; otherwise, unfolded protein aggregates are formed. Our study provided an example of a modified ribosome as a nexus for the quality control mechanism of synthesized protein.

## Results

### METTL18 is a histidine τ-N-methyltransferase

To overview τ-*N*-histidine methylation in cells and the corresponding enzymes, we quantified τ-*N*-methylhistidine by mass spectrometry (MS). To distinguish the two types of histidine methylations, we performed multiple reaction monitoring (MRM) of digested amino acids to trace specific *m/z* transitions from the precursor (Davydova et al., 2021) (see the “Materials and methods” section for details). Because SETD3 modifies abundant actin proteins (Dai et al., 2019; Guo et al., 2019; Kwiatkowski et al., 2018; Wilkinson et al., 2019; Zheng et al., 2020), *SETD3* knockout (KO) in HEK293T cells (Figure S1B-D) greatly reduced τ-*N*-methylhistidine (Figure 1A). However, a substantial fraction of τ-*N*- methylhistidine was left in the *SETD3* KO cells (Figure 1A), suggesting the presence of other mammalian τ-*N*-methyltransferase(s).

**Figure 1.**
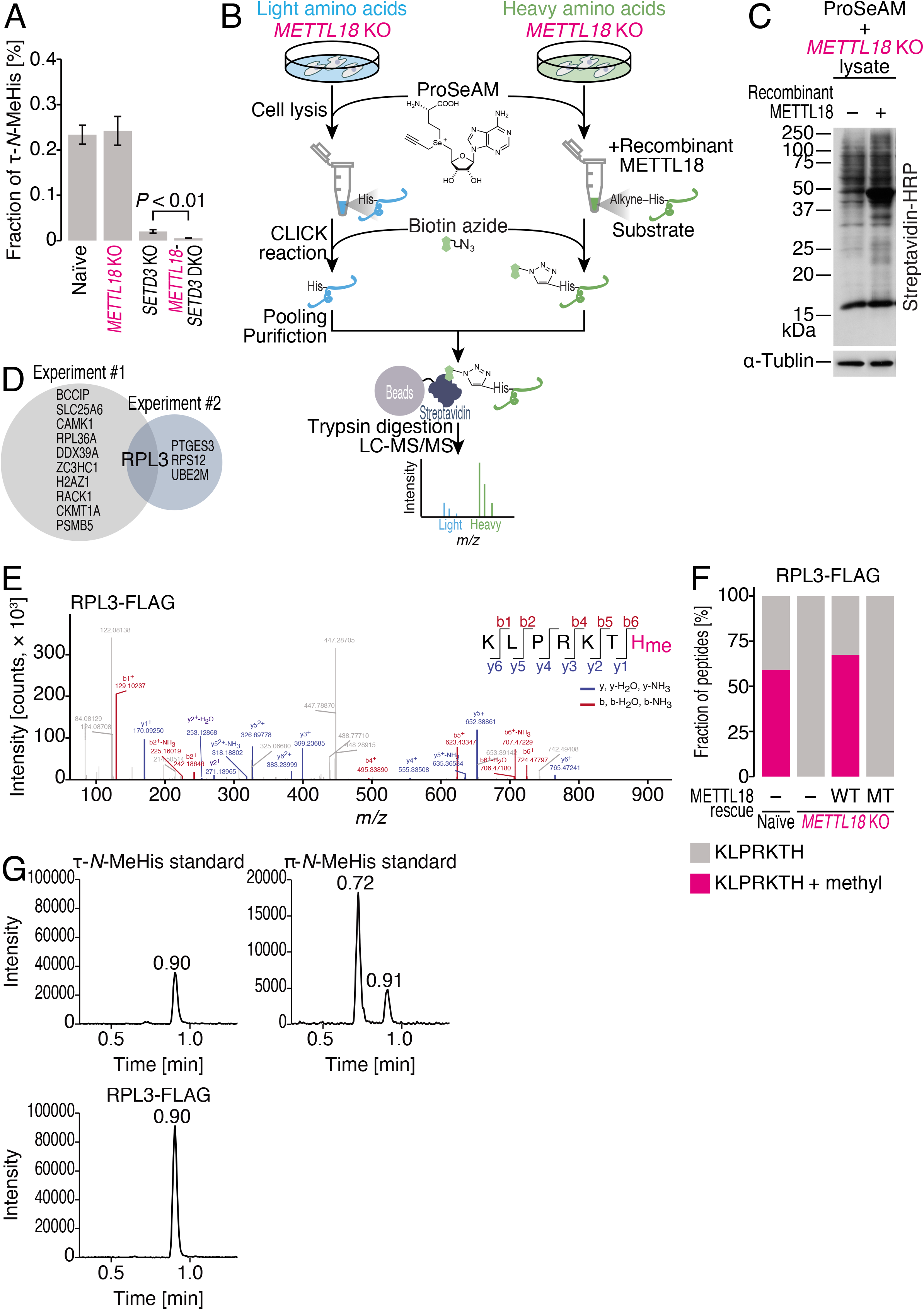
ProSeAM-SILAC identifies RPL3 as a substrate of METTL18. (A) MRM-based identification of τ-*N*-methylated histidine in bulk proteins from the indicated cell lines. Data represent the mean and s.d. (n =3). Significance was determined by Student’s *t*-test (unpaired, two sided). (B) Schematic representation of the ProSeAM-SILAC approach. (C) ProSeAM-labeled proteins in cell lysate with recombinant His-METTL18 protein. Biotinylated proteins were detected by streptavidin-HRP. Western blot for α-tubulin was used as a loading control. (D) Venn diagram of proteins identified in two independent experiments of ProSeAM-SILAC. The reproducibly detected protein was RPL3. (E) Methylated histidine residue in ectopically expressed RPL3-FLAG was searched by LC-MS/MS. (F) Quantification of methylated and unmethylated peptides (KLPRKTH) from the indicated cells. RPL3-FLAG was ectopically expressed and immunopurified for LC-MS/MS. (G) MRM-based identification of τ-*N*-methylhistidine in peptides from RPL3. The τ-*N*- methylhistidine standard, π*-N*-methylhistidine standard, and RPL3-FLAG peptide (KLPRKTH) results are shown. MeHis, methylhistidine. See also Figure S1 and S2.

An apparent candidate for the remaining τ-*N*-methyltransferase in *SETD3* KO cells was METTL18, whose yeast homolog (histidine protein methyltransferase 1, Hpm1) has been reported to catalyze τ-*N*-methylation on histidine in Rpl3 (Al-Hadid et al., 2014, 2016; Webb et al., 2010). Indeed, the *SETD3-METTL18* double-KO (DKO) cells (Figure S1B-D) had even lower levels of τ-*N*- methylhistidine than the *SETD3* single-KO cells (Figure 1A). Note that we could not detect any significant alteration in π*-N-* methylhistidine in any cells tested in this study (Figure S1E).

### METTL18 catalyzes τ-N-methylation on His245 in RPL3

These observations led us to survey the methylation substrate of METTL18. For this purpose, we harnessed propargylic *Se*-adenosyl-L-selenomethionine (ProSeAM), an analog of *S*-adenosyl-L-methionine (SAM). This compound acts as a donor in the methylation reaction by the SET domain and seven-β-strand methyltransferase (Davydova et al., 2021; Shimazu et al., 2014, 2018). Instead of a methyl moiety, a propargyl unit was added to the substrate residue, allowing biotin tagging with a click reaction (Davydova et al., 2021; Shimazu et al., 2014, 2018) (Figure 1B). Using ProSeAM, we performed *in vitro* methylation with recombinant METTL18 (Figure S1F) in the lysate of *METTL18* KO cells and detected the ProSeAM-reacted proteins in a METTL18 protein-dependent manner (Figure 1C). The high background signals are likely to have originated from methyltransferases other than METTL18 in the lysate.

For the quantitative and sensitive proteomic identification of methylated protein(s), we combined this approach with stable isotope labeling using amino acids in cell culture (SILAC) (Figure 1B). ProSeAM was added to the cell lysates of *METTL18* KO cells labeled with either light or heavy isotopic amino acids. Only the extract from heavy-isotope-labeled cells was incubated with recombinant METTL18 (Figure 1B). After lysate pooling and streptavidin purification (Figure 1B), isolated proteins were quantitatively assessed by liquid chromatography (LC)-MS/MS. Among the detected proteins, human RPL3 was the only candidate to be reproducibly identified (Figure 1D).

To ensure cellular histidine methylation on the protein, FLAG-tagged RPL3 was expressed in naïve HEK293T cells, immunopurified, and subjected to LC-MS/MS. This analysis identified the methylhistidine in cellular RPL3 and precisely annotated the residue at His245 (Figure 1E). In stark contrast, the same experiments with *METTL18* KO did not detect methylated His245 (Figure 1F). The ectopic expression of wild-type (WT) METTL18 in the KO cells rescued the modification, whereas mutations in the potential SAM binding site (Asp193Lys-Gly195Arg-Gly197Arg), which were predicted based on lysine methyltransferase orthologs (Ng et al., 2002), abolished the potential (Figure 1F). The same METTL18-dependent His245 methylation was also found in endogenous RPL3 (Figure S2C and S2D) isolated in the 60S subunit (Figure S2A and S2B).

To distinguish the methylation forms on histidine, we applied MRM to the peptide fragments containing His245 and clearly observed that methylation occurs at the τ-*N* position, but not at π*-N* position (Figure 1G).

Thus, taken together, our data demonstrated that His245 of RPL3 is a METTL18 substrate for the formation of τ-*N*-methylhistidine in cells.

### METTL18 associates with early pre-60S

As RPL3 in crude lysate was methylated *in vitro* (Figure 1D), we set out to recapitulate this reaction by purified factors using ^14^C-labeled SAM as a methyl donor. Irrespective of the human or mouse homolog, the immunopurified FLAG-tagged RPL3 proteins transiently expressed in *METTL18* KO cells were efficiently labeled by recombinant METTL18 (Figure 2A). Moreover, changing His245 to Ala completely abolished the reaction, validating His245 as a methylation site.

**Figure 2.**
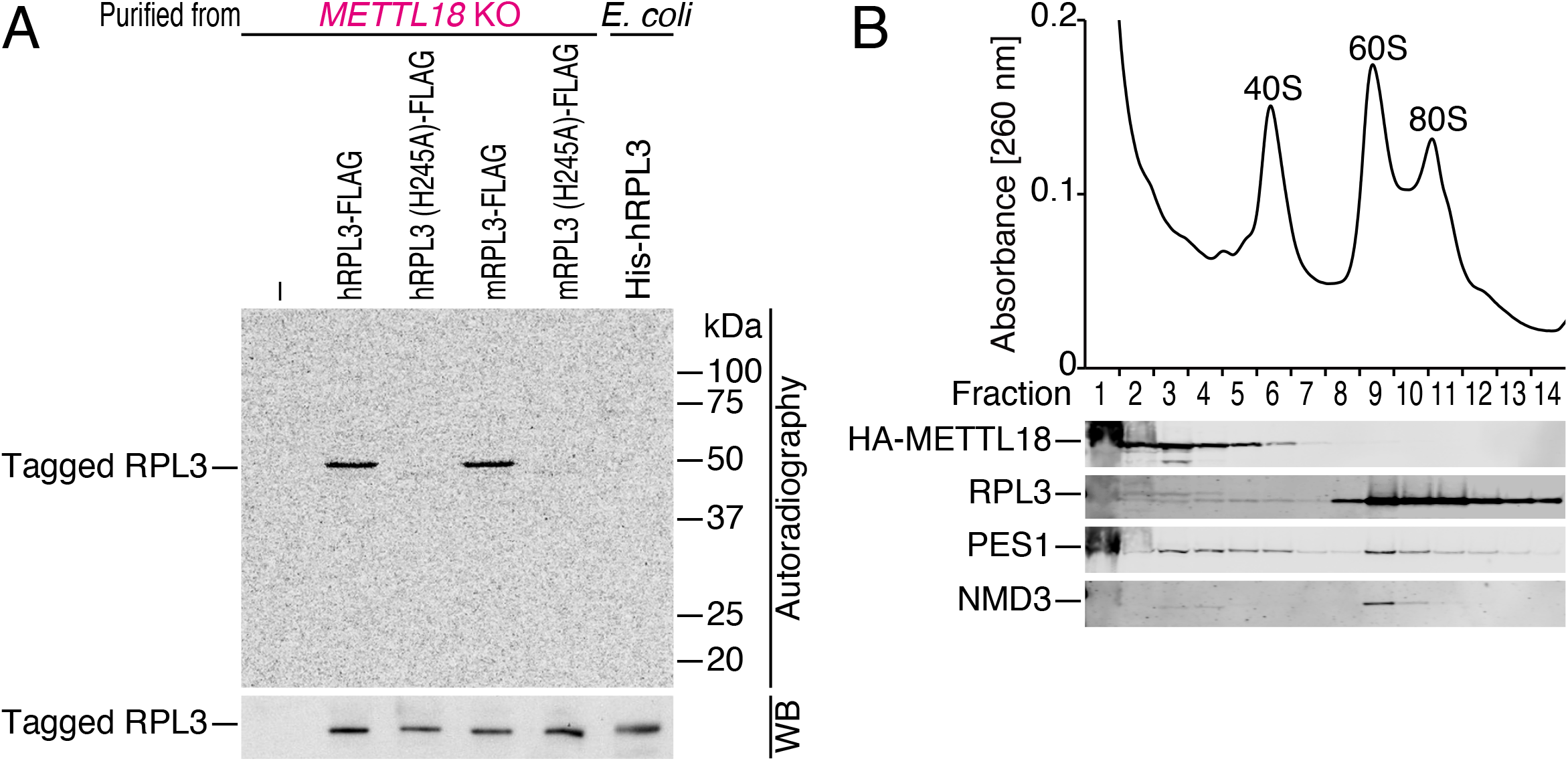
METTL18 associates with pre-60S. (A) *In vitro* methylation assay with recombinant His-GST-METTL18 protein and ^14^C-labeled SAM. Immunopurified human or mouse RPL3 expressed in *METTL18* KO cells and recombinant human RPL3 expressed in bacteria were used as substrates. (B) Western blot for the indicated proteins in ribosomal complexes separated by sucrose density gradient. See also Figure S3.

In contrast, recombinant RPL3 expressed in bacteria was a poor substrate (Figure 2A). A similar deficiency in the *in vitro* methylation assay by Hpm1 on solo RPL3 protein produced in bacteria was seen in yeast (Al-Hadid et al., 2014). Given that RPL3 expressed in mammalian cells assembled into ribosomes but RPL3 expressed in bacteria did not, we hypothesized that METTL18 recognizes RPL3 within ribosomes or preassembled intermediates. To characterize the molecular complex associated with METTL18, we separated the ribosomal complexes through a sucrose density gradient and found that METTL18 was associated with a complex smaller than mature 60S (Figure 2B). Given the smaller size, we speculated that the METTL18-associating complex is pre-60S in the middle of biogenesis. Indeed, the METTL18-containing subfractions also possessed PES1 (a yeast Nop7 homolog), a ribosome biogenesis factor at an early step (Kater et al., 2017; Sanghai et al., 2018), and a small portion of RPL3 (Figure 2B). On the other hand, NMD3, an adaptor protein of the late stage pre-60S for cytoplasmic export (Ma et al., 2017; Malyutin et al., 2017), was almost exclusive to the METTL18-associating pre-60S. These data suggested that RPL3, in an early intermediate complex of 60S biogenesis, is an efficient substrate for METTL18.

### Structural comparison of methylated and unmethylated His245 of RPL3 in ribosomes

The presence of METTL18 in the pre-60S complex led us to investigate the role of methylation in ribosome biogenesis. However, assessed by bulk 28S rRNA abundance (Figure S3A) and 60S fraction in sucrose density gradient (Figure S3B), no altered balance of the 60S subunit (relative to the 40S subunit) was observed in *METTL18* KO cells.

Thus, we hypothesized that RPL3 methylation may impact protein synthesis. To understand the potential role of methylation in RPL3, we reanalyzed published cryo-electron microscopy data (Osterman et al., 2020) and assessed the τ-*N*-methyl moiety on His245 of RPL3 (Figure 3A), which indicated that τ-*N*-methylated RPL3 is a stoichiometric component of the ribosome. A similar density of methylation on the histidine of RPL3 has also been found in rabbit ribosomes (Bhatt et al., 2021). On the other hand, the ribosomes isolated from *METTL18* KO cells lost density at the corresponding positions (Figure 3B, S4, and Table S1). The absence of methylation at the τ-*N* position allowed nitrogen to form hydrogen bonds with G1595 of 28S rRNA (Figure 3B). Given that G1595 is located in the loop of helix 35, which macrolide antibiotics target in bacterial systems (Kannan and Mankin, 2011), this difference in the interaction between His245 and G1595 suggests an alteration in the translation reaction.

**Figure 3.**
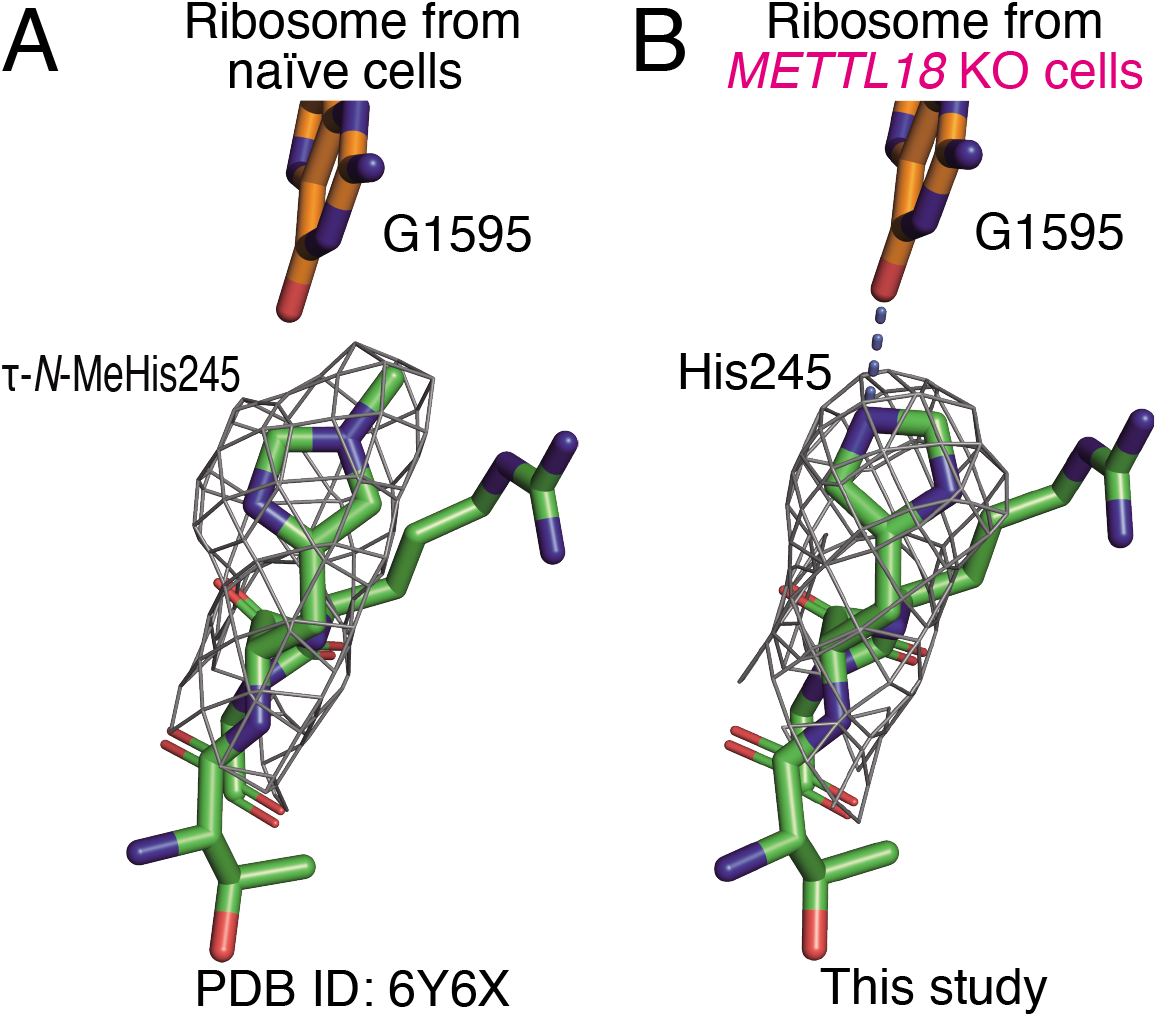
Structural differences in ribosomes upon methylation at His245. (A) Stick models of ^244^GHR^246^ of RPL3 and G1595 of the 28S rRNA of the human ribosome are shown with the cryo-EM density map around His245. The τ-*N*-methyl group was manually added to the original model (PDB ID: 6Y6X) (Osterman et al., 2020) based on the cryo-EM density map. (B) The same model as in (A) of human ribosome from *METTL18* KO cells. A hydrogen bond between His245 and G1595 is indicated with a dotted blue line. See also Figure S4 and Table S1.

### Methylation of His245 of RPL3 slows ribosome traverse at tyrosine codons

To investigate the impacts of the modification on protein synthesis, we assessed global translation in *METTL18* KO cells. However, we could not detect a significant difference in overall translation probed by polysome formation (Figure S5A) or nascent peptide labeling with *O*-propargyl-puromycin (OP-puro) (Figure S5B).

Nonetheless, we investigated the implications of RPL3 methylation defects across the transcriptome by ribosome profiling (Ingolia et al., 2009; Iwasaki and Ingolia, 2017). Strikingly, we found increased translation elongation of Tyr codons in *METTL18* KO cells; ribosome occupancy on Tyr codons at the A site was selectively reduced in the mutant cells (Figure 4A and 4B). This trend in ribosome occupancy was not observed at the P and E sites (Figure S6A and S6B). To evaluate the amino acid context associated with the high elongation rate, we surveyed the motifs around the A site with reduced ribosome occupancy by METTL18 depletion (Figure 4C) and analyzed the enriched/depleted sequence in the group (Figure 4D). Remarkably, Tyr at the A site was the predominant determinant for fast elongation in *METTL18* KO cells (Figure 4D).

**Figure 4.**
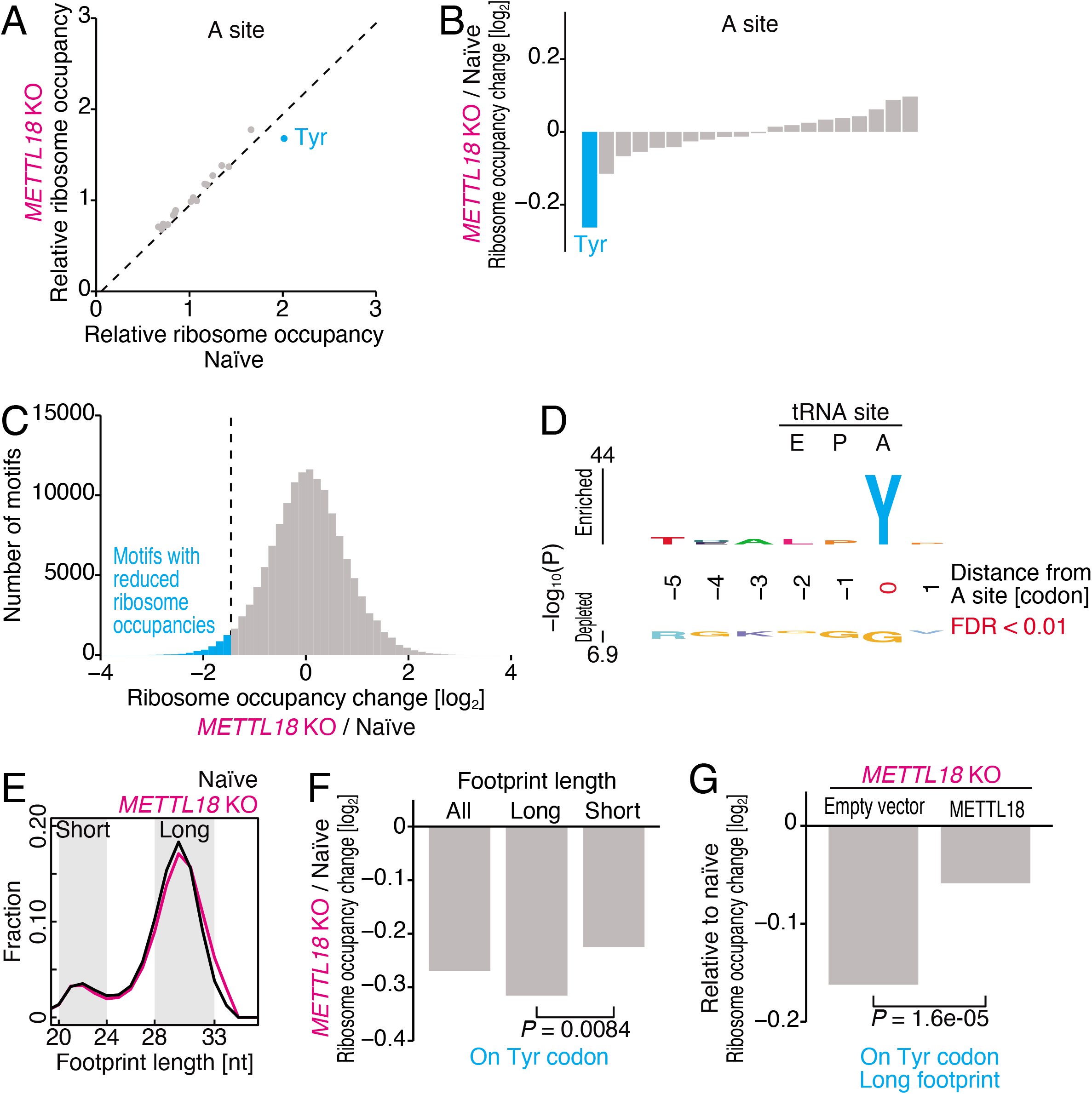
Ribosome profiling reveals Tyr codon-specific translation retardation by RPL3 methylation. (A) Ribosome occupancy at A-site codons in naïve HEK293T cells and *METTL18* KO cells. Data were aggregated into codons with each amino acid species. (B) Ribosome occupancy changes at A-site codons caused by METTL18 KO. (C) Histogram of ribosome occupancy changes in *METTL18* KO cells across motifs around A-site codons (7 amino acid motifs). Cyan: motifs with reduced ribosome occupancy (defined by ≤ mean – 2 s.d.). (D) Amino acid motifs associated with reduced ribosome occupancy in *METTL18* KO cells (defined in C) are shown relative to the A site (at the 0 position). (E) Distribution of footprint length in naïve HEK293T cells and *METTL18* KO cells. (F) Ribosome occupancy changes on Tyr codons by *METTL18* KO along all, long (28-33 nt), and short (20-24 nt) footprints. Significance was determined by Mann–Whitney *U* test. (G) The recovery of long footprint reduction in *METTL18* KO cells by ectopic expression of METTL18 protein. Significance was determined by Mann–Whitney *U* test. See also Figures S5 and S6.

Given the relatively long distance between the τ-*N*-methylated histidine and the mRNA codon, we reasoned that translocation rather than codon decoding is impacted by RPL3 methylation. Indeed, ribosome profiling data supported this scenario. Ribosome footprints possess two distinct populations of different length (short, peaked at ∼22 nt; long, peaked at ∼29 nt), reflecting the presence of A-site tRNA (Lareau et al., 2014; Wu et al., 2019) (Figure 4E). Since A-site tRNA-free ribosomes are more susceptible to RNase treatment at the 3′ end, the trimmed short footprints represent nonrotated ribosomes waiting for A site codon decoding by tRNA. On the other hand, long footprints originate from ribosomes accommodated with A-site tRNA in the middle of the peptidyl transfer reaction or subsequent subunit rotation (Lareau et al., 2014; Wu et al., 2019). The reduced footprints on Tyr codons in *METTL18* KO cells were more prominent in long footprints than short footprints (Figure 4F), suggesting that peptidyl transfer/subunit rotation rather than codon decoding is promoted in KO cells. Moreover, the rescue of the reduced long footprints on Tyr codons by ectopic expression of wild-type METTL18 further supported this conclusion (Figure 4G).

The smoother elongation may be caused by increased tRNA abundance. However, tRNA^Tyr^_GUA_, which decodes both UAU and UAC codons, was rather reduced by METTL18 depletion (Figure S6C). On the other hand, the abundance control tRNA^Leu^_HAG_ was not altered (Figure S6D). Thus, tRNA abundance did not explain the reduced footprints on Tyr codons.

These data together indicated the RPL3 methylation-mediated slowdown of translocation on the Tyr codons.

### RPL3 histidine methylation ensures the proper proteostasis

Ribosome traverse along the mRNA determines the quality of protein synthesized (Cassaignau et al., 2020; Collart and Weiss, 2019; Stein and Frydman, 2019). Slowdown of ribosome elongation is advantageous since it allows the duration of nascent protein folding before completion of protein synthesis. Therefore, we reasoned that translation elongation slowdown at Tyr codons by RPL3 methylation facilitates protein folding on ribosomes and maintains proper homeostasis of the proteome.

To test this possibility, we employed an aggregation-prone firefly luciferase (Fluc) reporter (Arg188Gln-Arg261Gln double mutant or DM) fused to enhanced green fluorescent protein (EGFP) (Gupta et al., 2011). Since this engineered protein necessitates chaperones to fold properly, the reduced pool of available chaperones (*i.e.*, proteotoxicity) leads to aggregation puncta of the reporter protein in cells. Indeed, METTL18 depletion induced the aggregation of FlucDM but not Fluc wild-type (WT) reporter (Figure 5A and 5B). As seen by the recovery of protein abundance by treatment with the proteasome inhibitor MG132 (Figure 5C), FlucDM protein was synthesized in *METTL18* KO cells of low quality and subjected to degradation.

**Figure 5.**
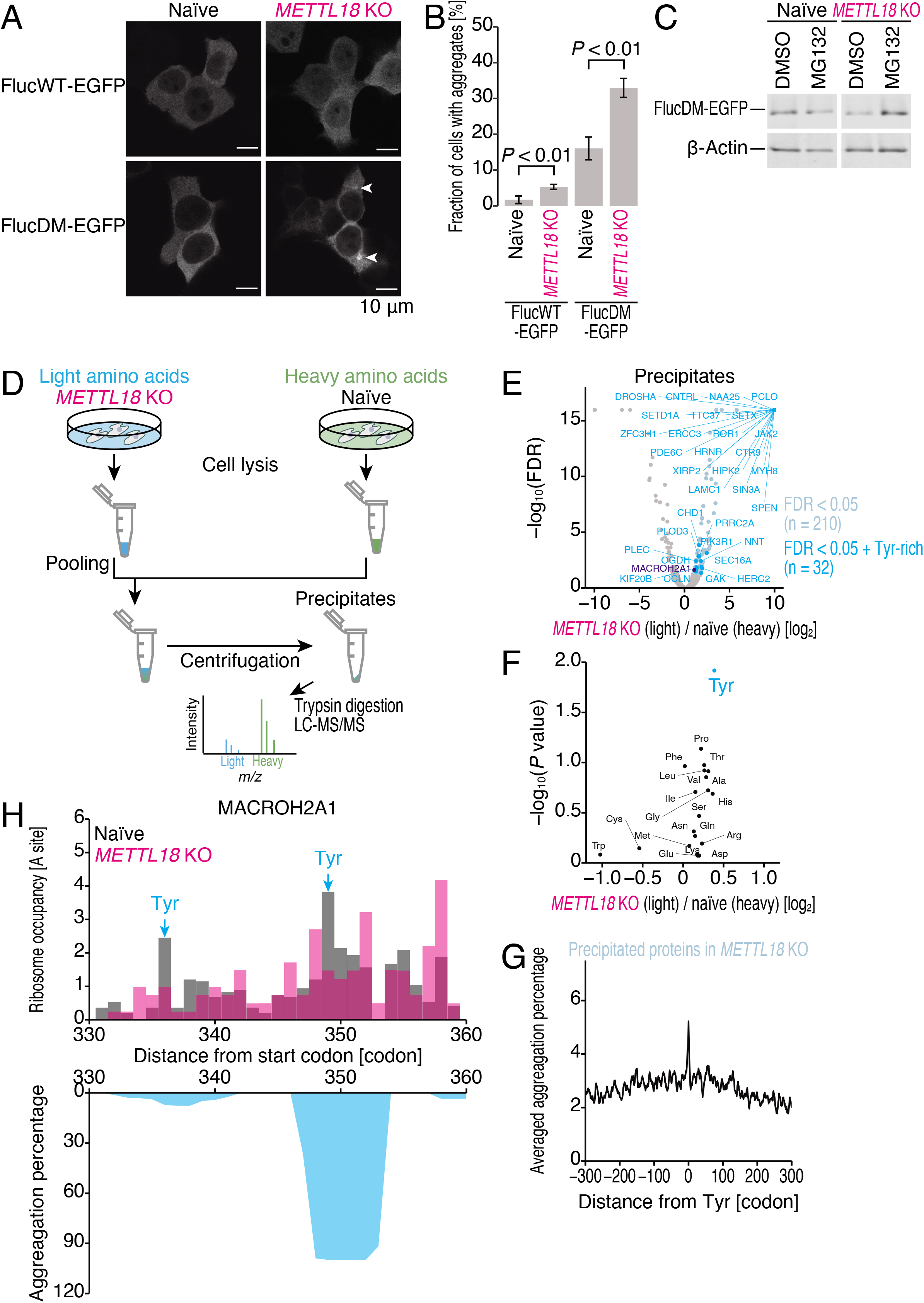
METTL18 deletion leads to cellular proteotoxicity. (A) Microscopic images of FlucWT-EGFP or FlucDM-EGFP in naïve HEK293T and *METTL18* KO cells. Scale bars are 10 μm. (B) Quantification of cells with Fluc-EGFP aggregates. Data present mean and s.d. (n = 3). Significance was determined by Student’s *t*-test (unpaired, two sided). (C) Western blot for FlucDM-EGFP (probed by anti-GFP antibody) expressed in naïve HEK293T and *METTL18* KO cells treated with MG132 (0.25 μM for 24 h). (D) Schematic representation of SILAC MS for precipitated proteins. (E) Volcano plot for precipitated proteins in *METTL18* KO cells, assessed by SILAC MS. Tyr-rich proteins were defined as proteins with 30 or more Tyr residues. (F) Amino acids associated with protein precipitation in *METTL18* KO cells. Precipitated proteins enriched with each amino acid were compared to the total precipitated proteome. The mean fold change and the significance (Mann-Whitney *U*-test) were plotted. (G) Metagene plot for aggregation percentage around Tyr codons of precipitated proteins in *METTL18* KO cells (defined in [E]). (H) Distribution (at the A site) of ribosome footprint occupancy along the MACROH2A1 gene in naïve HEK293T (gray) and *METTL18* KO (magenta) cells, depicted with the aggregation percentage (light blue) calculated with TANGO (Fernandez-Escamilla et al., 2004). Tyr codon positions are highlighted with arrows. See also Figures S7 and S8.

Then, we explored the proteome, the quality of which was assisted by RPL3 histidine methylation. For this purpose, we surveyed the proteins aggregated in *METTL18* KO cells by SILAC (Figure 5D). We observed that a subset of proteins were enriched in precipitates of *METTL18* KO cell lysates (Figure 5E). This subgroup significantly accumulated Tyr-rich proteins (defined as proteins possessing 30 Tyr or more) (hypergeometric test, *P* = 0.0068) (Figure 5E). The high probability of protein precipitates could not be explained by the increased net protein synthesis measured by ribosome profiling (Figure S7A). More generally, the Tyr-rich proteins were more prone to be precipitated by the deletion of METTL18 (Figure S7B) among all the amino acids (Figure 5F). Thus, proteomic analysis of cellular precipitates revealed that modulation of Tyr-specific translation elongation by RPL3 methylation confers proteome integrity.

These data led us to further investigate the properties of the proteome associated with Tyr. Here, we surveyed the aggregation propensity of the precipitated proteins by TANGO, which is based on statistical mechanics (Fernandez-Escamilla et al., 2004). Strikingly, meta-gene analysis showed the prominent enrichment of the TANGO-predicted aggregation propensity around the Tyr codons (Figure 5G). As exemplified in the MACROH2A1 protein, the subpart of the aggregation-prone region in this protein was found on Tyr with reduced ribosome occupancies in *METTL18* KO cells (Figure 5H). The correspondence among the aggregation-prone character of the protein, the cellular protein precipitates, and reduced ribosome occupancy indicates that Tyr translation modulated by RPL3-methylated ribosomes is associated with the quality of the synthesized proteins, preventing unwanted protein aggregation (Figure 6).

**Figure 6.**
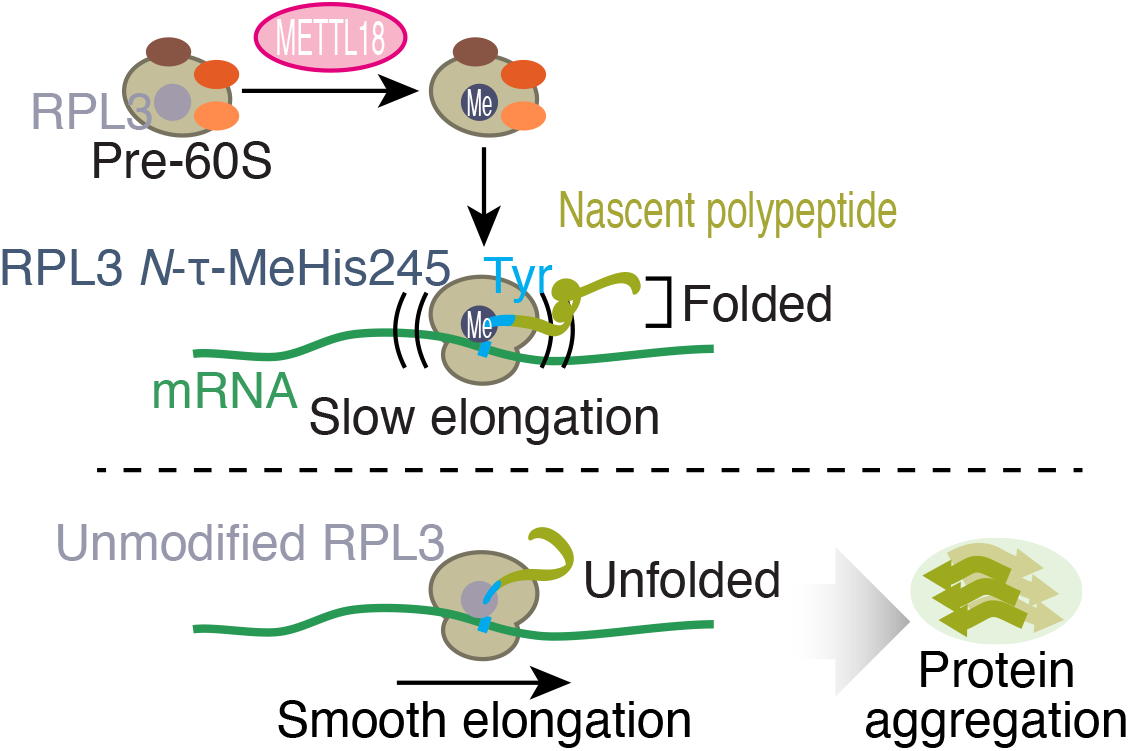
Schematic representation of METTL18-mediated control of translation and proteostasis. METTL18 adds a methyl moiety at the τ-*N* position of His245 in RPL3 in the form of an early 60S biogenesis intermediate. Methylated ribosomes slow the translocation of Tyr codons and extend the duration of nascent peptide folding, ensuring proteostatic integrity. Without RPL3 methylation, the accumulation of unfolded and ultimately aggregated proteins in cells was induced.

## Discussion

In addition to SETD3 (Dai et al., 2019; Guo et al., 2019; Kwiatkowski et al., 2018; Wilkinson et al., 2019; Zheng et al., 2020), METTL18 provides a second example of τ-*N*-methyltransferase. Whereas τ-*N*-methylation at the equivalent residue (His243) in yeast Rpl3 has been indirectly demonstrated (Webb et al., 2010), this study provided solid evidence (such as MS and cryo-EM) of the modification present on the homolog in humans.

Similar to prior yeast work (Al-Hadid et al., 2014), human RPL3 alone could not be methylated by METTL18. Rather, RPL3 was most likely to be modified in the early 60S biogenesis intermediate complex. This suggests that METTL18 recognizes a unique interface formed by RPL3 and other ribosome proteins and/or assembly factors. In the recent structural studies on pre-60 assembly intermediates (Kater et al., 2017; Sanghai et al., 2018), His245 of RPL3 in the early pre-60S in state B (Kater et al., 2017) is exposed to solvent (Figure S8A), possibly enabling access to METTL18, although His245 *per se* was not visible in the structure because of flexibility. In contrast, ribosomal proteins and rRNAs in the later stage of 60S biogenesis (at state D) (Kater et al., 2017) fill such a space (Figure S8B). Thus, histidine methylation by METTL18 should be restricted to specific timing of macromolecule construction.

In the course of this work, a paper from Falnes and coworkers was published and reached a similar conclusion regarding the methylation site (His245 in RPL3) and the methylation type (τ-*N* position) mediated by human METTL18 (Małecki et al., 2021). As we observed here, they also could not find alterations in the steady-state level of ribosome subunits in *METTL18* KO cells, although the slowdown of pre-rRNA cleavage was detected (Małecki et al., 2021), as was the case in yeast (Al-Hadid et al., 2016). Although the report showed that recombinant METTL18 protein methylated isolated ribosomes from *METTL18* KO, the data supported their observation that METTL18 localized in nucleoli (Al-Hadid et al., 2016) where ribosomes are still under assembly. These data could be interpreted as the purified ribosome may consist of contaminated pre-60S, and this fraction was an efficient substrate of METTL18.

The impact of METTL18 loss on translation differed in two reports. In the earlier report, ribosome profiling in *METTL18* KO cells revealed widespread effects of translation elongation, exemplified by slowdown of the GAA codon (Al-Hadid et al., 2016). The same experiment in this work showed the Tyr codon-specific enhancement of ribosome traverse (Figure 4). The difference may originate from the cells used (HAP1 haploid cells in the other report vs. HEK293T in this study). We believe that our condition may be a suitable setup to deduce the influence of METTL18 defects, since global protein synthesis was barely affected in our HEK293T cells (Figure S3A and S3B), while gene KO in HAP1 cells reduced the overall translation (Małecki et al., 2021).

The modification-mediated ribosome slowdown revealed in this study provides a unique example of translational control, since modification in general often offers a smooth translation, as exemplified in tRNA modifications (Nedialkova and Leidel, 2015; Tuorto et al., 2018). The exact mechanism of how His245 methylation in RPL3 leads to Tyr-specific elongation retardation remains elusive. In bacteria, the loop of helix 35, where unmethylated His245 interacts, was heavily modified (Kannan and Mankin, 2011), although the role of rRNA modification has been unclear. Eukaryotic RPL3 may functionally replace RNA modification by histidine methylation. From another viewpoint, His245 is located in the “basic thumb” region (Arg234-Arg246), which bridges the interaction of 28S rRNA helices (H61, H73, and H90) and facilitates the formation of a so-called “aminoacyl-tRNA accommodation corridor” (Meskauskas and Dinman, 2010). Thus, the methylation of His245 may allosterically alter the dynamic character of the aminoacyl-tRNA accommodation corridor to restrict the movement of charged Tyr on tRNA. Indeed, conformational changes in the aminoacyl-tRNA accommodation corridor were suggested by molecular dynamics simulation (Gulay et al., 2017).

We also did not exclude the possibility that this modification may impact other processes of translation. Indeed, an earlier report in yeast suggested that Hpm1 (a homolog of METTL18) deletion decreased the fidelity of translation, inducing stop codon readthrough (Al-Hadid et al., 2014, 2016). However, our ribosome profiling data in HEK293T cells did not show any increase in ribosome footprints in the 3′ UTR (Figure S8C and S8D) that could be accumulated by the stop codon readthrough (Arribere et al., 2016; Dunn et al., 2013). Analysis of the biochemistry, translatome, and proteome in a variety of organisms will reveal the generalized role of methylated RPL3 in protein synthesis.

Although ribosome traversal along the CDS is generally defined by the decoding rate in bacteria (Mohammad et al., 2019) and yeasts (Hussmann et al., 2015; Wu et al., 2019), the codon/amino acid-specific effects of RPL3 histidine methylation in humans suggest that the landscape is more complex in higher eukaryotes. Indeed, ribosome occupancy in mammals is poorly predicted by tRNA abundance in cells (Han et al., 2020). Thus, it is not surprising that interrogation of a wide array of ribosome modification codes, including RPL3 histidine modification, ultimately defines the speed of ribosome movement and thus the quality of protein synthesized in cells.

## Acknowledgments

We are grateful to all the members of the Iwasaki, Shinkai, Dohmae, and Ito laboratories for constructive discussion, technical help, and critical reading of the manuscript. This study was supported by the Support Unit for Bio-Material Analysis from RIKEN CBS Research Resources Division for mass spectrometry, Sanger sequencing, and confocal microscopy, and RIKEN ACCC for by supercomputer HOKUSAI SailingShip. This study used the following materials kindly provided: PX330-B/B from Tetsuro Hirose, pL-CRISPR.EFS.tRFP from Benjamin Ebert, pCI-neo Fluc-EGFP and pCI-neo FlucDM-EGFP from Franz-Ulrich Hartl, and a plasmid for *Salmonella* MTAN expression from Vern Schramm. S.I. was supported by a Grant-in-Aid for Transformative Research Areas (B) “Parametric Translation” (JP20H05784) from the Ministry of Education, Culture, Sports, Science and Technology (MEXT), a Grant-in-Aid for Young Scientists (A) (JP17H04998) and a Challenging Research (Exploratory) (JP19K22406) from the Japan Society for the Promotion of Science (JSPS), AMED-CREST (JP21gm1410001) from the Japan Agency for Medical Research and Development (AMED), and the Pioneering project (“Biology of Intracellular Environments”) and Aging Project from RIKEN. Yoichi S. was supported by a Grant-in-Aid for Scientific Research (A) (JP18H03991) and Scientific Research on Innovative Areas “Chromatin potential for gene regulation” (JP18H05530) from JSPS and the Pioneering project (“Epigenome manipulation”) from RIKEN. E.M.S. was supported by a Grant-in-Aid for Scientific Research (C) (JP21K06026) from JPSP and the Collaboration seed fund from RIKEN. Tadahiro S. was supported by a Grant-in-Aid for Scientific Research (C) (JP20K06497) from JSPS. T.I. was supported by a Grant-in-Aid for Scientific Research (B) (JP19H03172) by JSPS, the BDR Structural Cell Biology Project, the Pioneering Projects (“Dynamic Structural Biology” and “Biology of Intracellular Environments’’) and Aging Project from RIKEN, and AMED-CREST (JP21gm1410001) from AMED. DNA libraries were sequenced by the Vincent J. Coates Genomics Sequencing Laboratory at UC Berkeley, supported by NIH S10 OD018174 Instrumentation Grant. Structural analysis was supported by the Platform Project for Supporting Drug Discovery and Life Science Research (Basis for Supporting Innovative Drug Discovery and Life Science Research [BINDS], JP21am0101082) from AMED.

## Author contributions

Conceptualization, E.M.S., Tadahiro S., T.I., Yoichi S., and S.I.;

Methodology, E.M.S., Tadahiro S., T.I., Yoichi S., and S.I.;

Formal analysis, E.M.S., Tadahiro S., M.T., K.K., Takehiro S., and S.I.;

Investigation, E.M.S., Tadahiro S., M.T., K.K., and Takehiro S.;

Resources, Yoshihiro S., M.A., and M.S.;

Writing – Original Draft, S.I.;

Writing – Review & Editing, E.M.S., Tadahiro S., M.T., K.K., Takehiro S., N.D., T.I., Yoichi S., and S.I.;

Visualization, E.M.S., Tadahiro S., T.I., and S.I.;

Supervision, Tadahiro S., M.S., N.D., T.I., Yoichi S., and S.I.;

Funding Acquisition, E.M.S., Tadahiro S., T.I., Yoichi S., and S.I.

## Materials and methods

### Plasmid construction

#### PX330-B/B-gMETTL18

To express two guide RNAs targeting to upstream and downstream regions of exon 2 of *METTL18* gene, DNA fragments containing 5′-TCTCTTTAGCAGCTTATACA-3′ and 5′-GGTTGTGGATCAGGTTTACT-3′ were cloned into PX330-B/B (Yamazaki et al., 2018) via the BbsI and BsaI sites, respectively.

#### pL-CRISPR.EFS.tRFP-gSETD3

To express a guide RNA targeting exon 6 of the *SETD3* gene, a DNA fragment containing 5′-AGCCATGGGAAACATCGCAC-3′ was cloned into pL-CRISPR.EFS.tRFP (Addgene#57819) (Heckl et al., 2014).

#### pET19b-mMETTL18

To express N-terminally His-tagged full-length mouse METTL18, cDNA was PCR amplified from a FANTOM clone (AK139786) and cloned into NdeI and XhoI sites of the pET19b vector (Novagen).

#### pCold-GST-mMETTL18

To express N-terminally His- and GST-tagged full-length mouse METTL18, the PCR-amplified fragment was cloned into the NdeI and XhoI sites of the pCold-GST vector (TaKaRa).

#### pcDNA3-hRPL3-FLAG (WT and H245A) and pcDNA3-mRPL3-FLAG (WT and H245A)

To express C-terminally FLAG-tagged mouse and human RPL3, each cDNA fragment was amplified from the mouse cDNA and HEK293T cDNA libraries and cloned into EcoRI and NotI sites of the pcDNA3 vector (Invitrogen) with a C-terminal FLAG-tag sequence. To generate H245A mutants, the QuikChange Site-Directed Mutagenesis Kit (Agilent Technologies) was used.

#### pQCXIP-hMETTL18-HA and hMETTL18-Asp193Lys-Gly195Arg-Gly197Arg-HA

For the retrovirus expression, cDNA fragment of human METTL18 with a C-terminal HA sequence was cloned into AgeI and EcoRI sites of the pQCXIP vector (Clontech). To generate hMETTL18-Asp193Lys-Gly195Arg-Gly197Arg-HA, the QuikChange Site-Directed Mutagenesis Kit (Agilent Technologies) was used.

### Genome editing

KO cell lines were generated by the CRISPR-Cas9 system.

#### METTL18 KO cells

PX330-B/B-gMETTL18, which expresses hCas9 and two guide RNAs designed to induce large deletions in exon 2, and pEGFP-C1 (Clontech) were cotransfected into HEK293T cells. After 2 days of incubation, individual GFP-positive cells were sorted into 96-well plates. The clonal cell lines were screened by genomic PCR with the primers 5′-GGACTTTATGTTTGTCCAGGTGG-3′ and 5′-TGGGTTGTAAATGGTTTCTGAGG-3′.

#### METTL18 KO cells with stable METTL18 expression

Retrovirus packaging cells were transfected with pQCXIP-hMETTL18-HA or pQCXIP-hMETTL18-Asp193Lys-Gly195Arg-Gly197Arg-HA using PEI transfection reagent (Polysciences) and cultured for 24 h. Then, *METTL18* KO cells were inoculated with the virus-containing culture supernatant and 4 μg/ml of polybrene. At 24-h post infection, cells were selected with 1 μg/ml puromycin and cultured for additional 2 weeks.

#### SETD3 KO and SETD3-METTL18 DKO cells

pL-CRISPR.EFS.tRFP-gSETD3, which expressed a guide RNA designed to induce small insertion or deletion (InDel), was transfected into HEK293T cells or *METTL18* KO cells. After 2 days of incubation, individual RFP-positive cells were sorted into 96-well plates. The clonal cell lines were further screened by Western blot for SETD3 protein.

### Recombinant protein purification

#### Salmonella MTAN, His-METTL18, and His-GST-METTL18

BL21 (pLysS) strain transformed with *Salmonella* MTAN (Addgene, #64041), pET19b-mMETTL18, or pCold-GST-mMETTL18 was cultured in 2× YT medium with 100 µg/ml ampicillin and 0.2 mM isopropyl β-D-1-thiogalactopyranoside (IPTG) for 18 h at 16°C. The pelleted cells were lysed with 1× PBS with 0.5% NP-40 by sonication with a sonic homogenizer (Branson Ultrasonics, Sonifier S-250D) for 5 min on ice. After centrifugation at 15,000 × g for 10 min, the cleared cell extract was incubated with Ni-NTA Agarose (Qiagen) or Glutathione Sepharose 4B (Cytiva) for 1 h at 4°C with gentle agitation. The beads were washed 5 times with His wash buffer (50 mM Tris-HCl pH 7.4 and 25 mM imidazole) or GST wash buffer (1× phosphate-buffered saline [PBS]). Then, proteins were eluted with His elution buffer (50 mM Tris-HCl pH 7.4 and 250 mM imidazole) or GST elution buffer (50 mM Tris-HCl pH 8.0 and 50 mM glutathione). The purified proteins were dialyzed with dialysis buffer (50 mM Tris-HCl pH 8.0, 100 mM NaCl, 0.2 mM dithiothreitol [DTT], and 10% glycerol) using Slide-A-Lyzer Dialysis Cassettes (MWCO 10 kDa, Thermo Fisher Scientific) or Amicon Ultra (MWCO 10 kDa, Merck Millipore). The protein concentration was measured using the Bradford Protein Assay Kit (Bio-Rad).

### Western blot

Anti-α-tubulin (Sigma-Aldrich, clone B-5-1-2), anti-METTL18 (PROTEINTECH GROUP, 25553-1-AP), anti-SETD3 (Abcam, ab174662), anti-RPL3 (PROTEINTECH GROUP, 66130-1-lg and 11005-1-AP), anti-PES1 (Abcam, ab252849), anti-NMD3 (Abcam, ab170898), anti-HA (Medical & Biological Laboratories [MBL], M180-3), anti-GFP (Abcam, ab6556), and anti-β-actin (MBL, M177-3) primary antibodies were used.

For Figures 1C, 2A, and S1D, anti-mouse IgG, HRP-Linked Whole Ab Sheep (Cytiva, NA931V) and anti-rabbit IgG, HRP-Linked Whole Ab Donkey (Cytiva, NA934V) secondary antibodies were used. The chemiluminescence was raised with a Western Lightning Plus-ECL Kit (Perkin Elmer) according to the manufacturer’s protocol and detected with X-ray film (FUJI-FILM, RX-U). For biotinylated proteins, High Sensitivity Streptavidin-HRP (Thermo Fisher Scientific, 21130) was used.

To generate Figure 2B and 5C, IRDye680- or IRDye800CW-conjugated secondary antibodies (LI-COR Biosciences, 925-68070/71 and 926-32210/11, respectively) were used. Images were obtained with Odyssey CLx (LI-COR Biosciences).

### Mass spectrometry

#### MRM

Methylhistidine content analysis was performed essentially as previously described (Davydova et al., 2021). Proteins were precipitated with acetone and hydrolyzed to amino acids with 6 N HCl at 110°C for 24 h. After dissolving in 25 µl of 5 mM ammonium formate/0.001% formic acid, the amino acids were applied to a liquid chromatograph system (Thermo Fisher Scientific, Vanquish UHPLC). The amino acids loaded on a C18 column (YMC, YMC-Triart C18, 2.0 × 100 mm length, 1.9 µm particle size) were separated at a flow rate of 0.3 ml/min by gradient elution of mobile phase “A” (5 mM ammonium formate with 0.001% formic acid) and mobile phase “B” (acetonitrile) as follows: 0/0 – 1.5/0 – 2/95 – 4/95 – 4.1/0 – 7/0 (min/%B). The effluent was then directed to an electrospray ion source (Thermo Fisher Scientific, HESI-II) connected to a triple quadrupole mass spectrometer (Thermo Fisher Scientific, TSQ Vantage EMR) in positive ion multiple reaction monitoring mode. The electrospray was run with the following settings: spray voltage of 3000 V, vaporizer temperature of 450°C, sheath gas pressure of 50 arbitrary units, auxiliary gas pressure of 15 arbitrary units, and collision gas pressure of 1.0 mTorr. The transition of specific MH+→ fragment ions was monitored (His, *m/z* 156.1→83.3, 93.2, and 110.2; π*-N-*MeHis, *m/z* 170.1→95.3, 97.3, and 109.2; τ*-N-*MeHis, *m/z* 170.1→81.3, 83.3, and 124.2). Data were calibrated with 1 - 250 nM standards (His, π*-N-*MeHis, and τ*-N-*MeHis) at every run. The concentrations of His, π*-N-*MeHis, and τ*-N-*MeHis in the samples were calculated from the calibration curves obtained from the standards.

#### ProSeAM-SILAC-MS

ProSeAM was synthesized as previously described (Sohtome et al., 2018). ProSeAM substrate screening was carried out as reported (Shimazu et al., 2018) with some modifications. *METTL18* KO cells were cultured in Dulbecco’s modified Eagle’s medium (DMEM) containing either light Arg/Lys or heavy isotope-labeled Arg (^13^C_6_ ^15^N_4_ L-arginine)/Lys (^13^C_6_ ^15^N_2_ L-lysine) (Thermo Fisher Scientific, respectively) at least six doubling times. The cells were lysed with 50 mM Tris-HCl pH 8.0, 50 mM KCl, 10% glycerol, and 1% *n*-dodecyl-β-D-maltoside. For the cells cultured with the heavy amino acids, the lysate containing 200 μg of proteins was incubated with 150 μM ProSeAM and 10 μg of His-METTL18 in 50 mM Tris-HCl pH 8.0 at 20°C for 2 h. For the lysate with the light amino acids, His-METTL18 was omitted from the reaction. The reaction was stopped by the addition of 4 volumes of ice-cold acetone. The proteins were precipitated by centrifugation at 15,000 × g for 5 min, washed once with ice-cold acetone, and then dissolved in 58.5 µl of 1× PBS containing 0.2% sodium dodecyl sulfate (SDS). After the addition of 15 µl of 5× click reaction buffer (7.5 mM sodium ascorbate [Nacalai Tesque], 0.5 mM TBTA [AnaSpec], and 5 mM CuSO_4_) and 1.5 µl of 10 mM Azide-PEG4-Biotin (Click Chemistry Tools), the click reaction was conducted for 60 min at room temperature, stopped with 4 volumes of ice-cold acetone, and precipitated as described above. The protein in the pellet was resuspended in 75 µl of binding buffer (1× PBS, 0.1% Tween-20, 2% SDS, and 20 mM dithiothreitol [DTT]) and sonicated for 10 s. The light and heavy isotope-labeled samples were pooled in 450 µl of IP buffer (Tris-buffered saline [TBS] and 0.1% Tween-20) containing 3 µg of Dynabeads M-280 Streptavidin (Thermo Fisher Scientific) and incubated for 30 min at room temperature (note that the final SDS concentration in the solution was 0.5%). The protein-bound beads were washed three times with bead wash buffer (1× PBS, 0.1% Tween-20, and 0.5% SDS) and twice with 100 mM ammonium bicarbonate (ABC) and then used for Western blotting and MS.

For MS/MS analysis, the beads were incubated in 20 mM DTT and 100 mM ABC for 30 min at 56°C and then for 30 min at 37°C in the dark with supplementation with 30 mM iodoacetamide. Subsequently, proteins were digested with 1 µg trypsin (Promega) and subjected to liquid chromatography (Thermo Fisher Scientific, EASY-nLC 1000) coupled to a Q Exactive Hybrid Quadrupole-Orbitrap Mass Spectrometer (Thermo Fisher Scientific) with a nanospray ion source in positive mode, as previously reported (Davydova et al., 2021). The peptides loaded on a NANO-HPLC C18 capillary column (0.075-mm inner diameter × 150 mm length, 3 µm particle size, Nikkyo Technos) were eluted at a flow rate of 300 nl/min with the two different slopes of mobile phase “A” (water with 0.1% formic acid) and mobile phase “B” (acetonitrile with 0.1% formic acid): 0%–30% of phase B in 100 min and 30%–65% of phase B in 20 min. Subsequently, the mass spectrometer in the top-10 data-dependent scan mode was run with following parameters: spray voltage, 2.3 kV; capillary temperature, 275°C; mass-to-charge ratio, 350-1800; normalized collision energy, 28%. The MS and MS/MS data obtained with Xcalibur software (Thermo Fisher Scientific) were surveyed in the Swiss-Prot database with Proteome Discoverer (version 2.3, Thermo Fisher Scientific) and MASCOT search engine software (version. 2.7, Matrix Science). Peptides with false discovery rates (FDRs) less than 1% were considered. Proteins with the following criteria were defined as METTL18-dependent labeled proteins: 1.5-fold or more increase in heavy amino acid sample compared to light amino acid sample; one, 10% or more coverage of the protein; and 3 or more peptides identified.

#### LC-MS/MS for methylated peptide

The SDS-PAGE-separated and Coomassie staining-visualized proteins were excised and destained. The gel slices were reduced with 50 mM DTT and 4 M guanidine-HCl at 37°C for 2 h, followed by alkylation with 100 mM acrylamide at 25°C for 30 min. The RPL3 proteins were digested with chymotrypsin. The MS and MS/MS spectra were acquired with a Q Exactive HFX (Thermo Fisher Scientific). The mass spectrometer was operated in positive mode. The MS/MS spectra were obtained using a data-dependent Top 10 method. The acquired data were processed using Proteome Discoverer (version 2.3, Thermo Fisher Scientific). The processed data were used to search with MASCOT (version 2.7, Matrix Science) against the in-house database including the amino acid sequences of RPL3, using the following parameters: type of search, MS/MS ion search; enzyme, none; fixed modification, none; variable modifications, Gln->pyro-Glu (N-term Q), Oxidation (M), Propionamide (C), and Methyl (H); mass values, monoisotopic; peptide mass tolerance, ± 15 ppm; fragment mass tolerance, ± 30 mmu; peptide charge, 1+, 2+, and 3+; instrument type, ESI-TRAP.

#### SILAC-MS for precipitated proteins

Isotopically heavy amino acids (0.1 mg/ml ^13^C_6_^15^N_2_ L-lysine-HCl and 0.1 mg/ml ^13^C_6_^15^N_4_ L-arginine-HCl [both FUJIFILM Wako Chemicals]) or regular amino acids (0.1 mg/ml L-lysine-HCl and 0.1 mg/ml L-arginine-HCl [both FUJIFILM Wako Chemicals]) were added to DMEM deficient in both L-lysine and L-arginine for SILAC (Thermo Fisher Scientific) supplemented with 10% dialyzed fetal bovine serum (FBS) (Sigma-Aldrich). Naïve HEK293T cells and *METTL18* KO cells were cultured in media with heavy and light isotopes, respectively, for 2 weeks.

After a brief wash with PBS, cells were lysed with buffer containing 20 mM Tris-Cl pH 7.5, 150 mM KCl, 5 mM MgCl_2_, 1% Triton X-100, and 1 mM DTT. The protein concentration was measured using Qubit 2.0 Fluorometer (Thermo Fisher Scientific). The same amounts of proteins from light and heavy isotope-labeled lysates were mixed and centrifuged at 500 × g for 3 min. The supernatant was further centrifuged at 20000 × g and 4°C for 15 min. The precipitate was collected and used for analysis.

Filter-aided sample preparation (FASP) was used for protein digestion (Wiśniewski et al., 2009). The precipitate was dissolved in SDT lysis buffer (4% [w/v] SDS, 0.1 M DTT, and 100 mM Tris-HCl pH 7.6) and incubated at 95°C for 5 min. Denatured samples were diluted 10 times with UA buffer (8 M urea and 100 mM Tris-HCl pH 8.5) and loaded to Vivacon 500, 30,000 MWCO Hydrosart (Sartorius) to trap the protein on the filter. The filter unit was washed with the UA buffer. Proteins on the filter unit were alkylated with 100 μl of the IAA solution (0.05 M iodoacetamide in UA). Then, the filter unit was washed with 100 μl of UA buffer three times and 100 μl of 50 mM NH_4_HCO_3_ three times. Proteins on the filter unit were digested with 40 μl of trypsin solution (trypsin [V5111, Promega] in 50 mM NH_4_HCO_3_) at 37°C for 18 h. Then, the peptides were eluted from the filter unit by 40 μl of NH_4_HCO_3_ twice and 50 μl of 0.5 M NaCl once and then pooled.

LC-MS/MS analysis was performed using EASY-nLC 1000 (Thermo Fisher Scientific) and Q Exactive (Thermo Fisher Scientific) equipped with a nanospray ion source. The peptides were separated with a NANO-HPLC capillary column C18 (0.075 x 150 mm, 3 μm, Nikkyo Technos) at 300 nl/min flow rate with strep gradients of solvent A (0.1% formic acid) and solvent B (acetonitrile with 0.1% formic acid); 0%-30% B for 100 min and then 30%-65% B for 20 min. The resulting MS and MS/MS data were searched against the Swiss-Prot database using Proteome Discoverer (version 2.4, Thermo Fisher Scientific) with MASCOT search engine software (version 2.7, Matrix Science). The peptides with FDR of 0.05 or less were considered for the subsequent analysis. To quantify the difference of peptides labeled with differential isotopes, built-in SILAC 2-plex quantification method in Proteome Discoverer (version 2.4, Thermo Fisher Scientific) was used. Peptide abundances were normalized by total peptide amount. *P*-values were calculated by *t*-test (background-based) and then adjusted by Benjamini-Hochberg method.

### Methylation assay

FLAG-tagged RPL3 was transiently expressed in *METTL18* KO cells and immunopurified with ANTI-FLAG M2 Affinity Gel (Sigma-Aldrich). The purified proteins were incubated in 1× reaction buffer (50 mM Tris-HCl pH 8.5 and 50 mM MgCl_2_) with 1 µg of His-GST-METTL18, 2 µM MTAN, and 0.01 µCi of ^14^C-labeled SAM (Perkin Elmer) at 30°C for 2 h. The reaction was stopped by the addition of Laemmli SDS-sample buffer. Proteins were separated on a 10% acrylamide SDS-PAGE gel. The dried gel was exposed to an imaging plate (FUJI-FILM) for 48 h. The autoradiograph was detected with a phosphor imaging scanner (Cytiva, Amersham Typhoon).

### Cryo-EM

The crude ribosomal pellet was suspended in buffer A (50 mM Tris-HCl pH 7.5, 150 mM KCl, 4 mM magnesium acetate, 1 mM DTT, 7% [w/v] sucrose, and 1 mM puromycin) by mixing with a magnetic stirrer for 4 h on ice. After stirring, the suspension was centrifuged at 15,000 × g for 10 min at 4°C. The supernatant was loaded onto a HiPrep 16/60 Sephacryl S-500 column (Cytiva) equilibrated with buffer A without puromycin. The fraction was recovered and concentrated using Amicon Ultra (MWCO 50 kDa, Merck Millipore). The concentrated mixture was supplemented with 0.1 mM puromycin, 2 mM magnesium acetate, and 2 mM ATP and incubated for 30 min at 37°C. The reaction was loaded onto a 10–40% (w/v) sucrose gradient with buffer B (50 mM Tris-HCl pH 7.5, 500 mM KCl, 4 mM magnesium acetate, and 2 mM DTT) and centrifuged at 25,000 rpm in an SW28 rotor for 16 h at 4°C. Two-milliliter fractions were successively fractionated from the top of the gradient. An aliquot of each fraction was analyzed by SDS-PAGE, and the gels were stained with Coomassie Brilliant Blue (CBB) to detect the 40S and 60S subunit proteins. The 40S- and 60S-subunit fractions were mixed and concentrated using Amicon Ultra filter units (MWCO 50 kDa, Merck Millipore). The concentrated mixture was loaded onto a 10–50% (w/v) sucrose gradient with buffer C (50 mM Tris-HCl pH 7.5, 150 mM KCl, 10 mM magnesium acetate, and 2 mM DTT) and centrifuged at 28,000 rpm in an SW41Ti rotor for 3 h at 4°C. Gradient fractionation was carried out using a piston gradient fractionator equipped with a TRIAX flow cell detector (Biocomp) by continuous monitoring of absorbance at a wavelength of 280 nm. The fractions containing 80S ribosomes were dialyzed against preparation buffer (10 mM HEPES-KOH pH 7.5, 30 mM potassium acetate, 10 mM magnesium acetate, and 1 mM DTT) and concentrated using Amicon Ultra filter units (MWCO 50 kDa, Merck Millipore). The concentrated sample was quantified by measuring absorbance at a wavelength of 260 nm and flash-cooled with liquid nitrogen.

For cryo-EM sample preparation, Quantifoil R1.2/1.3 300 mesh copper grids (Quantifoil) were covered with an amorphous carbon layer prepared in-house. The thawed sample was diluted to 70 nM (an absorbance at 260 nm of 3.5) with preparation buffer, and 3 μl of the sample was applied onto grids at 4°C at 100% relative humidity using a Vitrobot Mark IV (FEI). After incubation for 30 s and blotting for 3 s, the grids were plunged into liquid ethane.

The cryo-EM dataset was collected with a Tecnai Arctica transmission electron microscope (FEI) operated at 200 kV using a K2 summit direct electron detector (Gatan) (0.97 Å/pixel). The 5,517 images collected were fractionated to 40 frames, with a total dose of ∼50 e^-^/Å^2^.

Processing of cryo-EM data was performed with RELION-3.1 (Zivanov et al., 2020). The movie frames were aligned with MotionCor2 (RELION’s own implementation), and the CTF parameters were estimated with CTFFIND-4.1 (Rohou and Grigorieff, 2015). Particles were automatically picked using a template-free Laplacian-of-Gaussian (LoG) filter (250–500 Å), and 381,227 particles were extracted with twofold binning. After 2D classification, 136,688 particles were selected and applied to 3D classification. A low-pass-filtered (40 Å) map of the human 80S ribosome (EMD-9701) (Yokoyama et al., 2019) was used as a reference map for 3D classification, and 118,470 particles were selected after this step. These particles were re-extracted without rescaling and used in 3D refinement, Bayesian polishing, CTF refinement, and then 3D refinement again. After these steps, focused refinement with a mask on the 60S subunit and postprocessing resulted in a resolution of 2.72 Å.

For molecular modeling, the model of the 60S subunit from the human 80S ribosome structure at 2.9 Å resolution (PDB: 6QZP) (Natchiar et al., 2017) was used as a starting model and manually fitted into the map using UCSF Chimera (Pettersen et al., 2004). Map sharpening and model refinement were performed in PHENIX (Adams et al., 2010), and the model was further refined manually with Coot (Emsley et al., 2010). The modified nucleotides of ribosomal RNAs were introduced based on quantitative mass spectrometry data (Taoka et al., 2018).

### Sucrose density gradient

For Figure 2B, cells were lysed with whole-cell lysis buffer (50 mM Tris-HCl pH 7.5, 150 mM NaCl, 5 mM MgCl_2_, 1% NP-40, 1 mM DTT, and 100 µg/ml cycloheximide). The whole-cell lysates were passed through a 21-gauge needle and incubated at 4°C for 15 min. For Figure S5A, cells were lysed with lysis buffer (20 mM Tris-Cl pH 7.5, 150 mM KCl, 5 mM MgCl_2_, 1% Triton X-100, 1 mM DTT, and 100 µg/ml cycloheximide) and cleared by centrifugation at 20,000 × g and 4°C for 10 min. Cell lysate containing 40 µg of total RNA was loaded onto a 10-50% sucrose gradient and ultracentrifuged at 35,300 rpm and 4°C for 2.5 h by a Himac CP80WX ultracentrifuge (Hitachi) with a P40ST rotor (Hitachi).

For Figure S2A and Figure S3B, cells were lysed with EDTA lysis buffer (20 mM Tris-Cl pH 7.5, 150 mM KCl, 5 mM EDTA, 1% Triton X-100, and 1 mM DTT) and centrifuged at 20,000 × g and 4°C for 10 min. The supernatant with 40 µg of total RNA was ultracentrifuged at 35,300 rpm and 4°C for 2.5 h (Figure S2A) or at 38,000 rpm and 4°C for 4 h (Figure S3B).

After ultracentrifugation, fractions were concentrated using Amicon Ultra filter units (MWCO 10 kDa, Millipore) and used for Western blot (Figure 2B) or MS analysis (Figure S2B).

### Newly synthesized protein labeling by OP-puro

Metabolic labeling of nascent proteins with OP-puro was performed as previously described (Iwasaki et al., 2019). Cells were treated with 20 µM OP-puro and incubated at 37°C for 30 min in a CO_2_ incubator. After washing with PBS, cells were lysed with buffer containing 20 mM Tris-HCl pH 7.5, 150 mM NaCl, 5 mM MgCl_2_, and 1% Triton X-100. Nascent polypeptides were labeled with IRdye800CW Azide (LI-COR Biosciences) with a Click-iT Cell Reaction Buffer Kit (Thermo Fisher Scientific). After free dye was removed by a G-25 column (Cytiva), labeled polypeptides were separated by SDS-PAGE. The gel was imaged by Odyssey CLx (LI-COR Biosciences) for the detection of nascent peptides with infrared at 800 nm. Then, total proteins were stained with CBB (FUJIFILM Wako Chemicals) and imaged with an infrared 700 nm signal. The gel area ranging from 17 kDa to 280 kDa was quantified by Image Studio (version 5.2, LI-COR Biosciences), and the nascent peptide signal was normalized to the total protein signal.

### Ribosome profiling

#### Library preparation

Ribosome profiling was conducted as previously described (McGlincy and Ingolia, 2017; Mito et al., 2020). Cells were cultured in DMEM, high glucose, GlutaMAX Supplement (Thermo Fisher Scientific) supplemented with 10% FBS (Sigma-Aldrich) at 37°C in a humidified atmosphere containing 5% CO_2_.

Cell lysates were treated with RNase I, and 17-34-nt protected RNA fragments were gel-excised. After the RNA fragments were ligated with preadenylated linkers, ribosomal RNAs were removed by a Ribo-Zero Gold rRNA Removal Kit (Human/Mouse/Rat) (Illumina, RZG1224). Then, the ligated RNA fragments were reverse-transcribed. The cDNAs were circularized by CircLigaseII (Lucigen) and PCR-amplified. The libraries were sequenced on a HiSeq4000 (Illumina).

#### Data analysis

After removal of the linker sequences, all reads were aligned to human noncoding RNAs (including rRNA, tRNA, snoRNA, snRNA, and microRNA) using STAR (version 2.7.0a) (Dobin et al., 2013). Then, the remaining reads were aligned to the human hg38 reference genome by STAR and assigned to canonical transcripts in the University of California, Santa Cruz (UCSC) known gene reference.

The A-site offsets for each footprint length were empirically estimated: in the data set with naïve HEK293T and *METTL18* KO cells, 15 for 20-22, 24, and 28-30 nt long footprints and 16 for 23 and 32-33 nt long footprints, in the data set with *METTL18* KO cells with METTL18 WT and empty vector expression, 15 for 28-31 nt long footprints and 16 for 32-33 nt long footprints. The reads of each codon were normalized by the average reads per codon of the transcript. We excluded the first and last 5 codons from the analysis. Transcripts with an average reads per codon of 0.3 or higher were considered. The value of ribosome occupancy was the average of normalized reads at each position.

To search motifs associated with Tyr codon around A site, ribosome occupancies on the 7 amino acid motifs were averaged. Motifs with average score less than 0.3 were excluded from the downstream analysis. Then the log2-fold changes in *METTL18* KO cells over naïve cells were calculated. Motifs with log2-fold changes less than mean – 2 s.d. were subjected to kpLog (http://kplogo.wi.mit.edu) (Wu and Bartel, 2017).

### Northern blot

Total RNA was extracted from cells by TRIzol Reagent (Thermo Fisher Scientific) according to the manufacturer’s instructions. Purified RNAs were electrophoresed on Super Sep RNA gels (FUJIFILM Wako Chemicals), transferred onto nylon membranes (Biodyne, Thermo Fisher Scientific), and then UV-crosslinked. The membranes were incubated with UltraHyb-Oligo (Thermo Fisher Scientific) at 37°C for 1 h. DNA oligonucleotide probes (see below for details) were radiolabeled with [γ−^32^P] ATP (PerkinElmer) by T4 PNK (New England Biolabs) and purified with a G-25 column (Cytiva). After prehybridization, membranes were incubated with a labeled DNA probe overnight at 37°C and washed three times with 2 × saline-sodium citrate (SSC) solution. The signal on the membranes was detected by an Amersham Typhoon (Cytiva) scanner. DNA oligonucleotide sequences used as probes are listed below. tRNA^Tyr^_GUA_: 5′-ACAGTCCTCCGCTCTACCAGCTGA-3′, tRNA^Leu^_HAG_: 5′-CAGCGCCTTAGACCGCTCGGCCA-3′, and U6: 5′-CACGAATTTGCGTGTCATCCTT-3′. The background-subtracted signal was quantified by ImageQuant TL (Cytiva).

### Proteotoxic-stress reporter assay

pCI-neo Fluc-EGFP (Addgene plasmid #90170; http://n2t.net/addgene:90170; RRID: Addgene_90170) or pCI-neo FlucDM-EGFP (Addgene plasmid #90172; http://n2t.net/addgene:90172; RRID: Addgene_90172) (kind gifts from Franz-Ulrich Hartl) was transfected into naïve cells or *METTL18* KO cells by *Trans*IT-293 (Mirus) according to the manufacturer’s instructions. For immunofluorescence staining, cells were cultured on a Nunc Lab-Tek II - CC^2^ Chamber Slide system (Thermo Fisher Scientific) and fixed with 4% paraformaldehyde in PBS for 20 min at room temperature. After being washed twice with PBS containing 0.2% Triton X-100, the cells were incubated with Intercept (TBS) Blocking Buffer (LI-COR Biosciences) with 0.2% Triton X-100 for 1 h at room temperature. After removal of the blocking buffer, the cells were incubated overnight at 4°C with anti-GFP antibody (Abcam, ab1218) diluted 1:1000 with Intercept (TBS) Blocking Buffer with 0.2% Triton X-100. Then, the cells were washed three times with PBS containing 0.2% Triton X-100 and incubated for 1 h at room temperature with anti-mouse secondary antibody conjugated with Alexa 488 (Thermo Fisher Scientific, R37120) diluted 1:1000 with Intercept (TBS) Blocking Buffer (LI-COR Biosciences) with 0.2% Triton X-100. Then, the cells were washed three times with PBS containing 0.2% Triton X-100. Slide chambers were mounted with VECTASHIELD HardSet Antifade Mounting Medium (VECTOR LABORATORIES). Immunofluorescence images were acquired by a FLUOVIEW FV3000 (Olympus).

To count the aggregation foci, 2 × 10^5^ cells were seeded in 35-mm glass-bottom dishes (IWAKI) and cultured for 24 h. Cells were transfected with 2 µg of pCI-neo Fluc-EGFP or pCI-neo FlucDM-EGFP with 4 µg of PEI (Polysciences) and cultured for an additional 24 h. GFP fluorescence was observed under an FLUOVIEW FV3000 (Olympus). At least one hundred GFP-positive cells were observed to count cells with aggregates (n=3).

For MG132 treatment, cells were treated with 0.25 µM MG132 at 24 h post transfection. After subsequent 24-h culture, cells were harvested as described in the “Ribosome profiling” section.

### Accession numbers

The ribosome profiling data obtained in this study (GSE179854) were deposited to the National Center for Biotechnology Information (NCBI). The proteome data for ProSeAM SILAC (ID: PXD026813) were available via ProteomeXchange. The structural coordinates (PDB: 7F5S) and cryo-EM maps (EMDB: EMD-31465) of His245 methylation-deficient ribosomes have been deposited in the Protein Data Bank (PDB) and Electron Microscopy Data Bank (EMDB), respectively.

---

**Figure S1.**
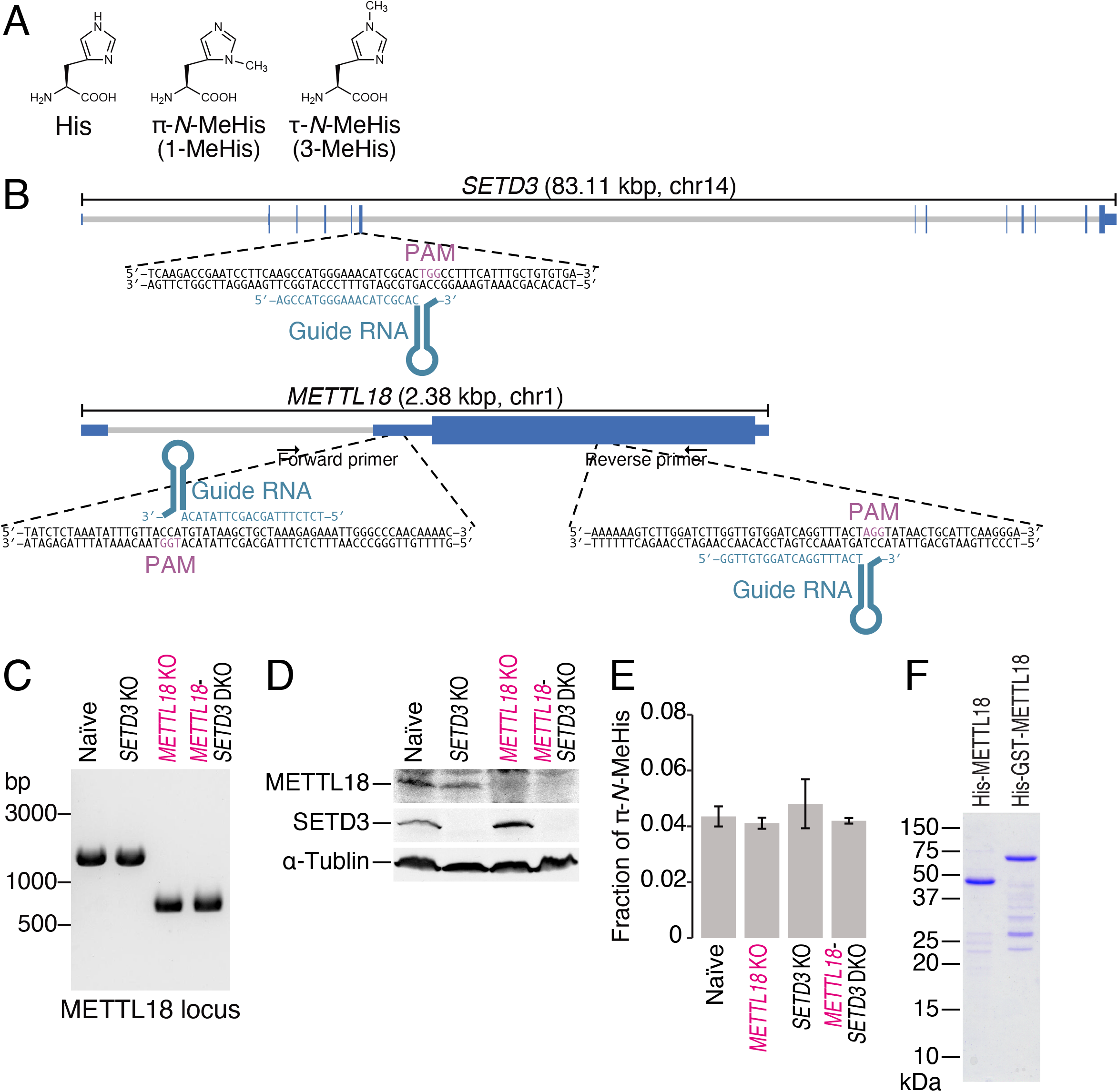
Generation of *SETD3* and *METTL18* KO cells; related to Figure 1. (A) Chemical structure of histidine, π*-N-*methylhistidine, and τ*-N-*methylhistidine. (B) Schematic representation of guide RNAs (gRNAs) designed for CRISPR-Cas9-mediated gene knockout. (C) Genomic PCR validated the partial deletion of the *METTL18* gene locus. (D) Western blot of the indicated proteins to confirm the knockout of *SETD3* and *METTL18*. (E) MRM-based identification of π-*N*-methylated histidine in bulk proteins from the indicated cell lines. Data represent the mean and s.d. (n =3). MeHis, methylhistidine. (F) Coomassie brilliant blue (CBB) staining of recombinant METTL18 proteins used in this study.

**Figure S2.**
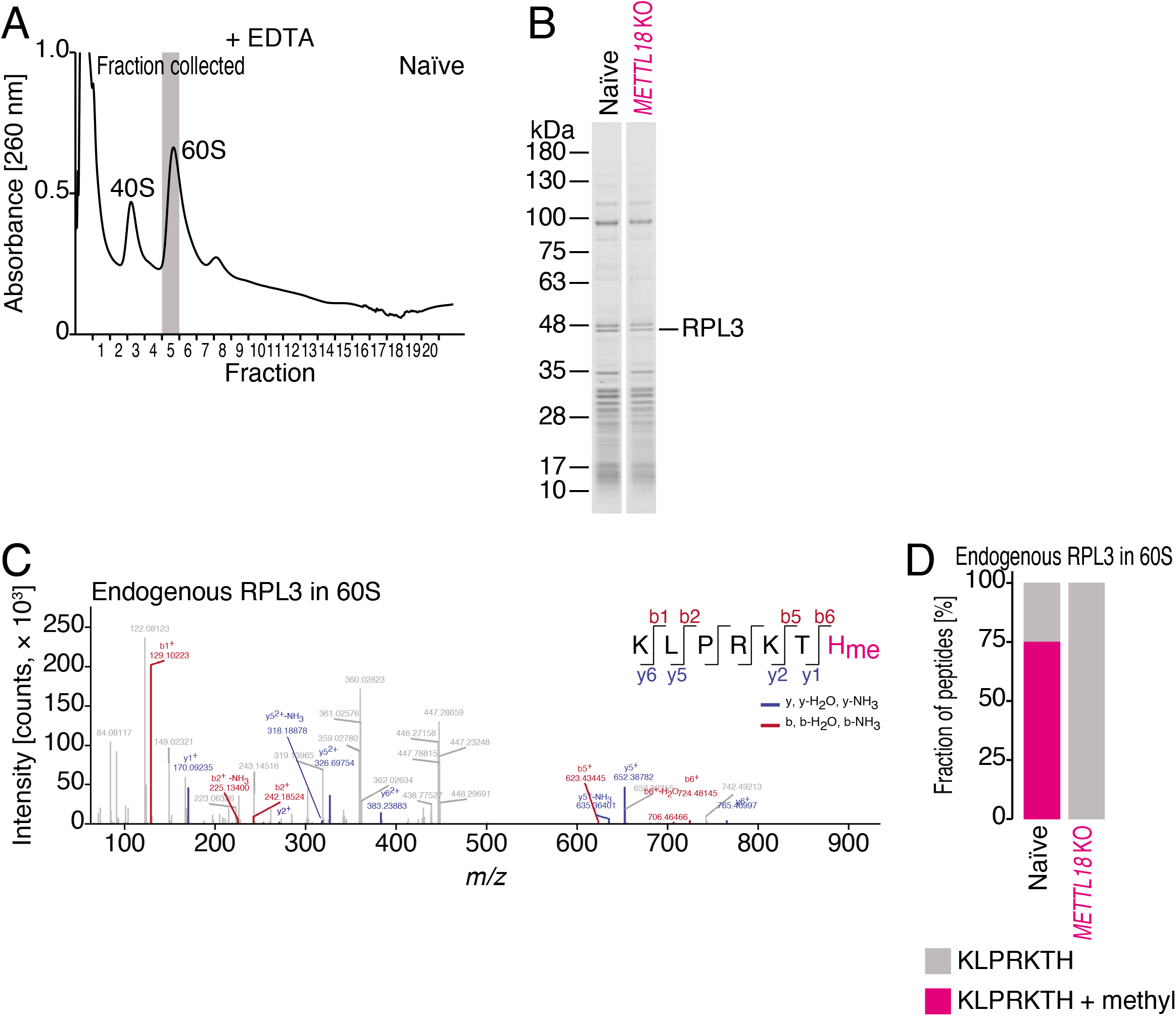
Characterization of methylhistidine in endogenous RPL3; related to Figure 1. (A) Sucrose density gradient for ribosomal complexes. Lysate was prepared with a buffer containing EDTA to dissociate 80S into 40S and 60S. The 60S fraction used for LC-MS/MS analysis is highlighted in gray. (B) CBB staining of proteins in the 60S fraction in naïve HEK293T and *METTL18* KO cells. (C) Methylated histidine residue in endogenous RPL3 in 60S was searched by LC-MS/MS. (D) Quantification of methylated and unmethylated peptide (KLPRKTH) from endogenous RPL3 in 60S cells.

**Figure S3.**
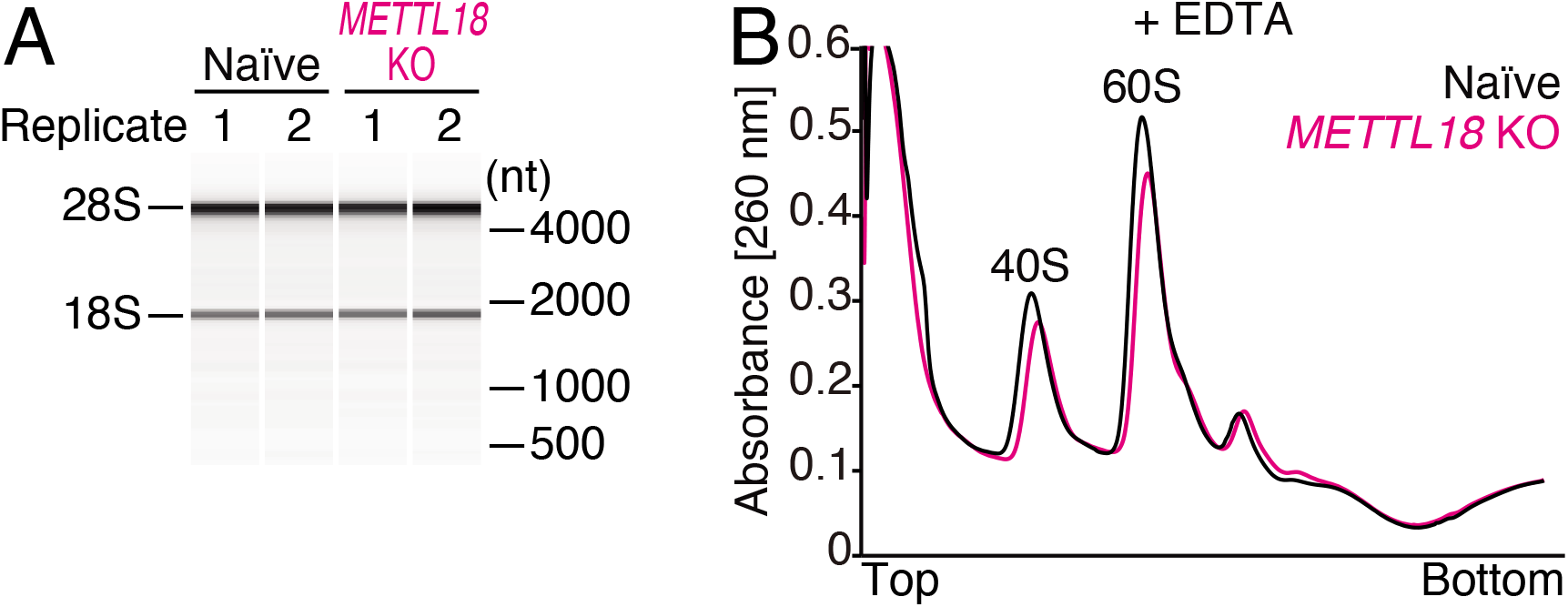
Ribosome subunit ratio in *METTL18* cells; related to Figure 3. (A) Electropherogram of ribosomal RNAs from naïve HEK293T and *METTL18* KO cells. (B) Sucrose density gradient for ribosomal complexes from naïve HEK293T and *METTL18* KO cells. The lysate was prepared with a buffer containing EDTA to dissociate 80S into 40S and 60S.

**Figure S4.**
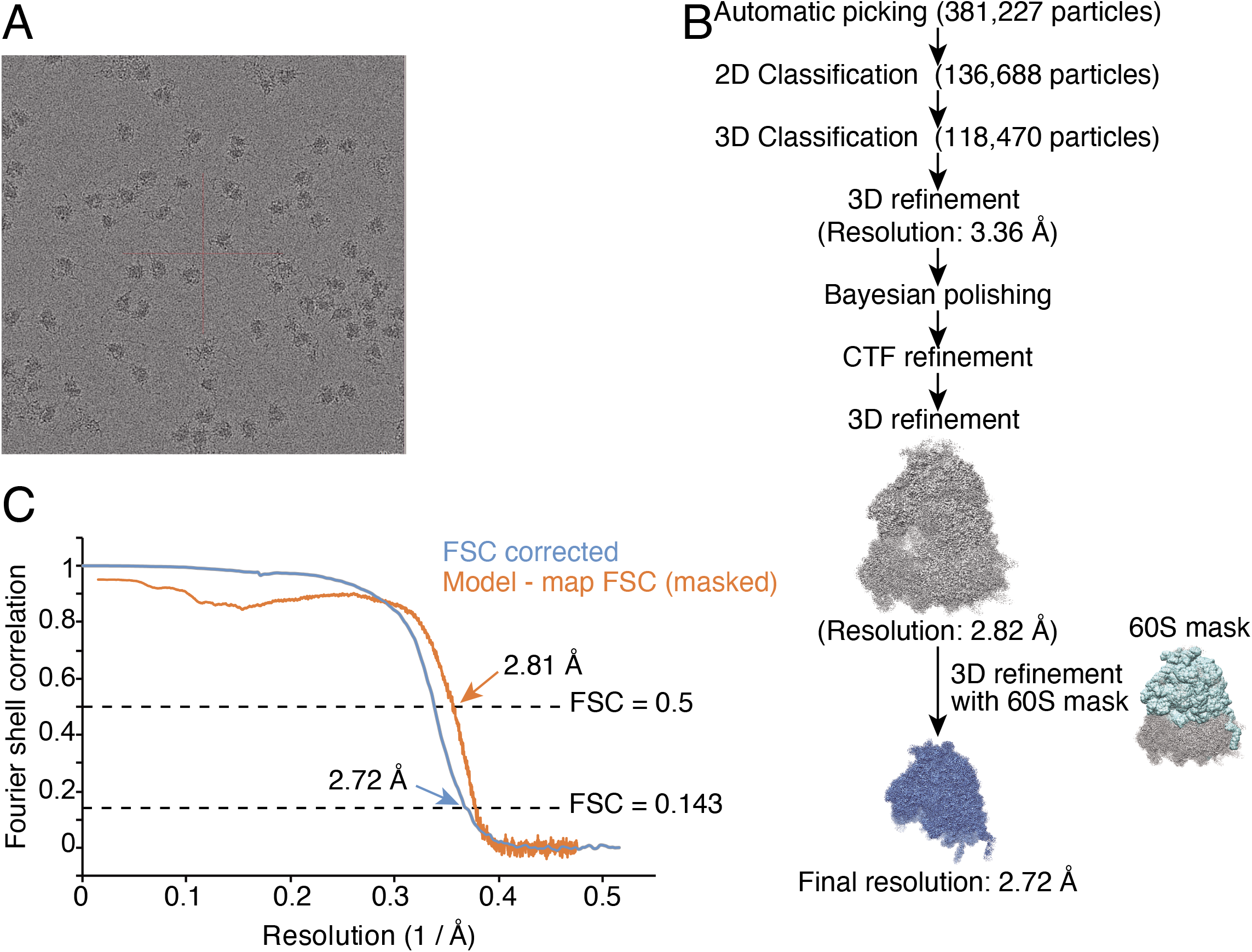
Characterization of the structure of the 60S subunit from *METTL18* KO cells; related to Figure 3. (A) Representative cryo-EM micrographs of human ribosomes isolated from *METTL18* KO cells. (B) Flow of the cryo-EM structural analysis of the human 60S subunit from *METTL18* KO cells. (C) Resolution curves of the reconstituted cryo-EM structure of the human 60S subunit from *METTL18* KO cells.

**Figure S5.**
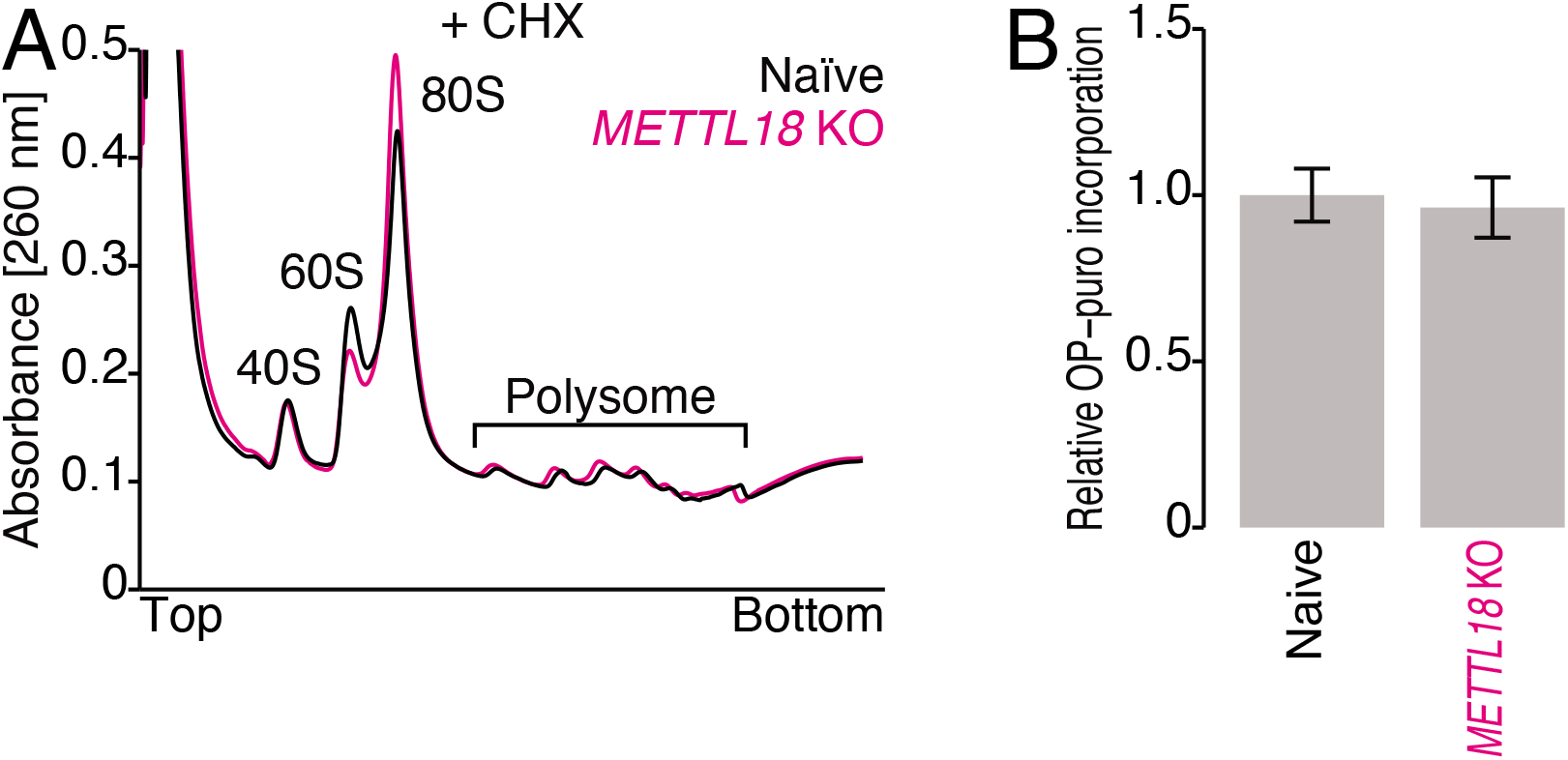
Basal translation activity in *METTL18* cells; related to Figure 4. (A) Sucrose density gradient for ribosomal complexes from naïve HEK293T and *METTL18* KO cells. 80S and polysomes were stabilized by Mg ion and cycloheximide. (B) Newly synthesized proteins in naïve HEK293T and *METTL18* KO cells were labeled with OP-puro and then conjugated with infrared 800 (IR800) dye with a click reaction. The signal was normalized to total proteins stained with CBB. Data present mean and s.d. (n =3).

**Figure S6.**
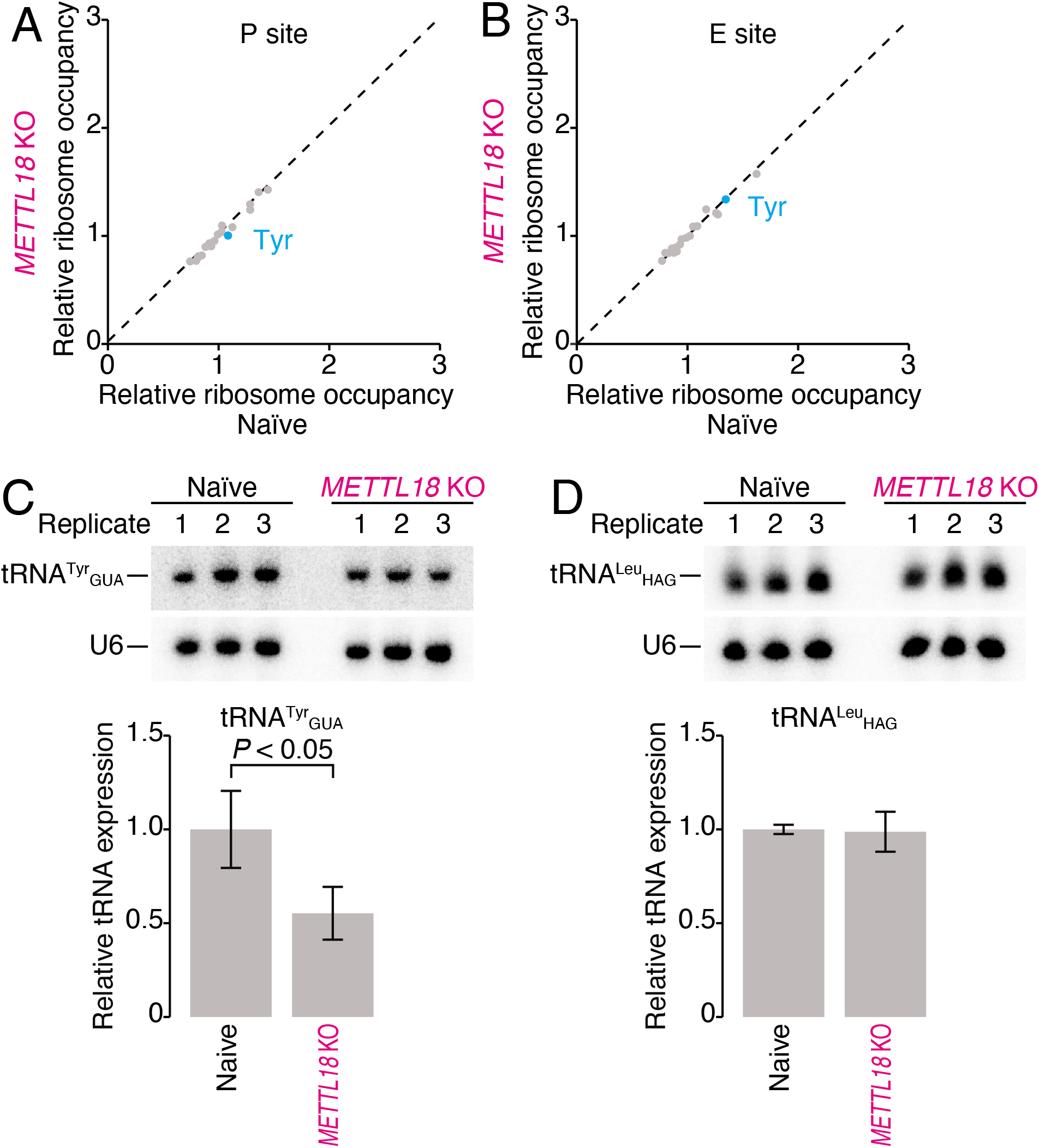
Characterization of ribosome occupancy monitored by ribosome profiling; related to Figure 4. (A and B) Ribosome occupancy at P-site (A) and E-site (B) codons. Data were aggregated into codons with each amino acid species. (C) Northern blot for tRNA^Tyr^_GUA_ (top) and its quantification (bottom). U6 snRNA was used as loading control. Data present mean and s.d. (n =3). Significance was determined by Student’s *t*-test (unpaired, two sided). (D) Same as (C) but for tRNA^Leu^_HAG_. H stands for A, C, or U.

**Figure S7.**
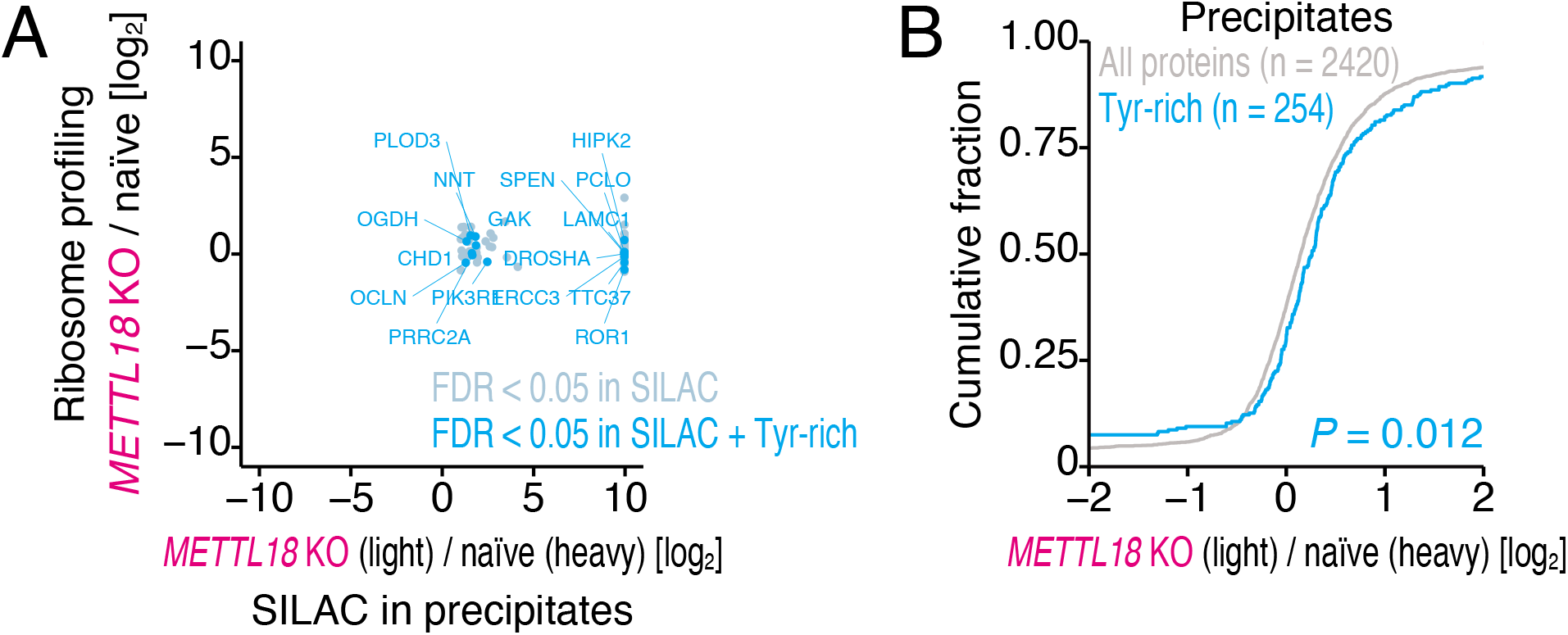
Characterization of precipitated proteins identified by SILAC mass spectrometry; related to Figure 5. (A) Comparison of fold change in protein precipitates assessed by SILAC and in ribosome footprints assessed by ribosome profiling by METTL18 depletion. (B) Cumulative distribution of Tyr-rich proteins along the fold change in protein precipitates by METTL18 depletion. Significance was calculated by Mann-Whitney *U*-test.

**Figure S8.**
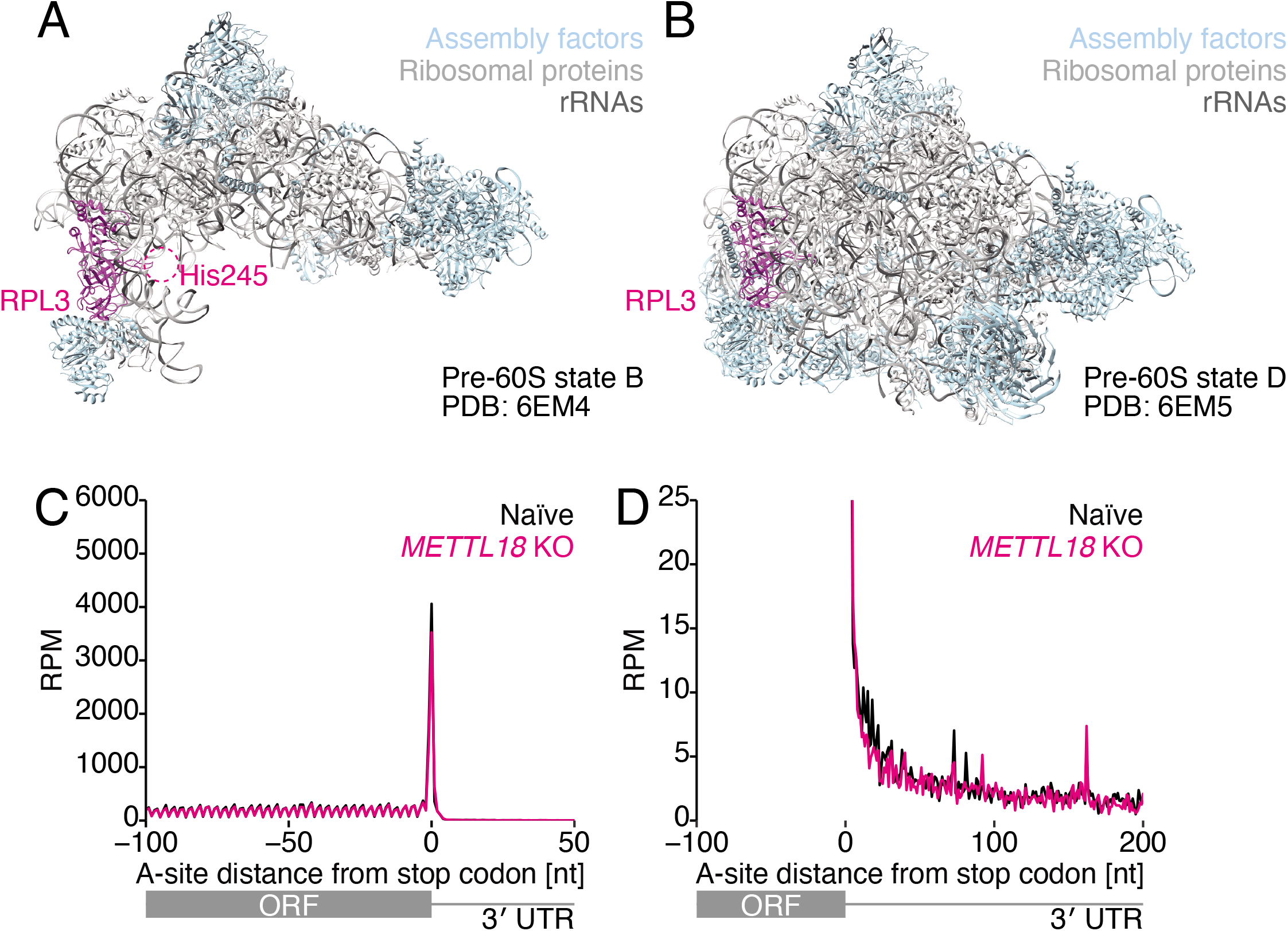
Comparison of the structure of the early and late pre-60S as a possible RPL3 methylation target; related to Figure 5. (A and B) Structures of early (state B, PDB 6EM4:) and late (state D, PDB: 6EM5) pre-60S (Kater et al., 2017). A possible region of the protein fragment containing His245 in RPL3 is highlighted in a dashed circle. RPL3, magenta; assembly factor, light blue; ribosomal proteins, light gray; rRNA, dark gray. (C and D) Metagene analysis of ribosome footprints around the stop codon. A-site potion of footprints is depicted. In D, a zoomed-in view of the plot is shown.

**Table S1.**
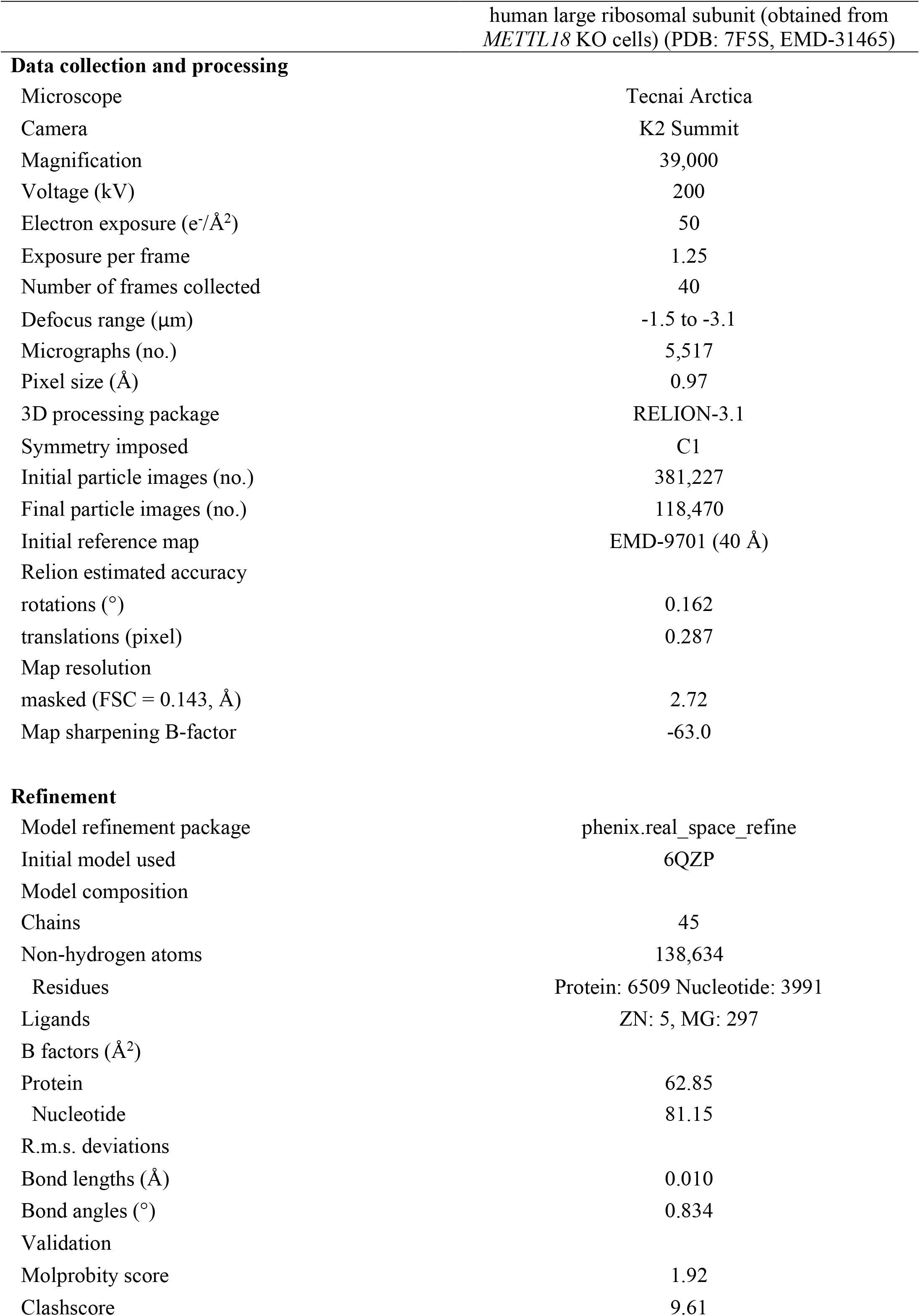
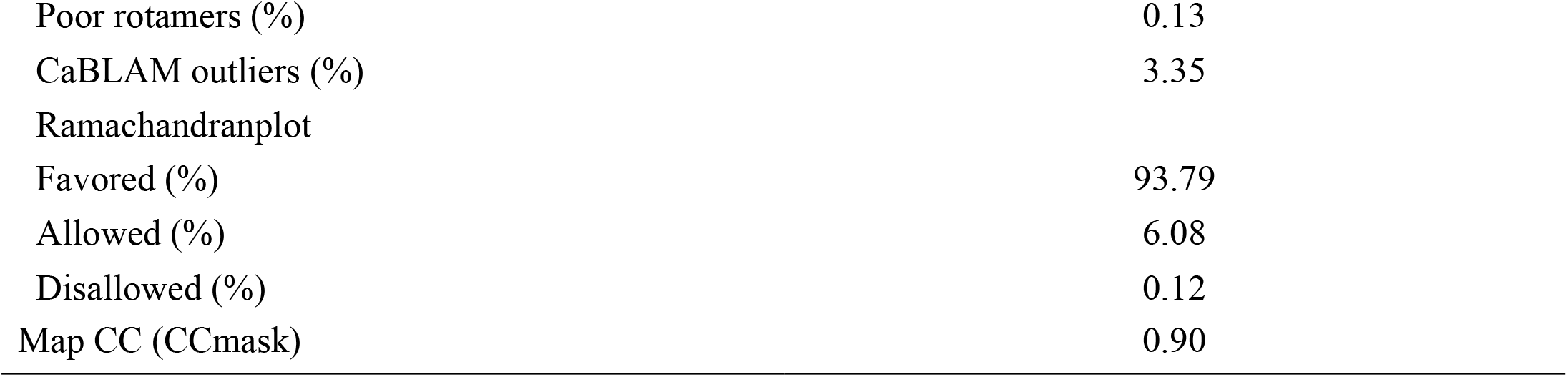
Data collection, model building, refinement, and validation statistics for cryo-EM data obtained in this study; related to Figure 3.

**Data collection and processing**
|  |  |
| --- | --- |
| Microscope | Tecnai Arctica |
| Camera | K2 Summit |
| Magnification | 39,000 |
| Voltage (kV) | 200 |
| Electron exposure (e<sup>-</sup>/Å<sup>2</sup>) | 50 |
| Exposure per frame | 1.25 |
| Number of frames collected | 40 |
| Defocus range (μm) | -1.5 to -3.1 |
| Micrographs (no.) | 5,517 |
| Pixel size (Å) | 0.97 |
| 3D processing package | RELION-3.1 |
| Symmetry imposed | C1 |
| Initial particle images (no.) | 381,227 |
| Final particle images (no.) | 118,470 |
| Initial reference map | EMD-9701 (40 Å) |
| Relion estimated accuracy |  |
| rotations (°) | 0.162 |
| translations (pixel) | 0.287 |
| Map resolution |  |
| masked (FSC = 0.143, Å) | 2.72 |
| Map sharpening B-factor | -63.0 |
| **Refinement** |  |
| Model refinement package | phenix.real_space_refine |
| Initial model used | 6QZP |
| Model composition |  |
| Chains | 45 |
| Non-hydrogen atoms | 138,634 |
| Residues | Protein: 6509 Nucleotide: 3991 |
| Ligands | ZN: 5, MG: 297 |
| B factors (Å<sup>2</sup>) |  |
| Protein | 62.85 |
| Nucleotide | 81.15 |
| R.m.s. deviations |  |
| Bond lengths (Å) | 0.010 |
| Bond angles (°) | 0.834 |
| Validation |  |
| Molprobity score | 1.92 |
| Clashscore | 9.61 |
| Poor rotamers (%) | 0.13 |
| CaBLAM outliers (%) | 3.35 |
| Ramachandranplot |  |
| Favored (%) | 93.79 |
| Allowed (%) | 6.08 |
| Disallowed (%) | 0.12 |
| Map CC (CCmask) | 0.90 |

